# Tunable dynamics in a multi-strain transcriptional pulse generator

**DOI:** 10.1101/2022.09.23.509237

**Authors:** David M. Zong, Mehdi Sadeghpour, Sara Molinari, Razan N. Alnahhas, Andrew J. Hirning, Charilaos Giannitsis, William Ott, Krešimir Josić, Matthew R. Bennett

## Abstract

A major challenge in synthetic biology is the manipulation of engineered gene circuits toward a specified behavior. This challenge becomes more difficult as synthetic systems become more complex by incorporating additional genes or strains. Here we demonstrate that circuit dynamics can be tuned in synthetic consortia through the manipulation of strain fractions within the community. To do this, we constructed a microbial consortium comprised of three strains of engineered *Escherichia coli* that, when co-cultured, use homoserine lactone (HSL) mediated intercellular signaling to create a multi-strain incoherent type-1 feedforward loop (I1-FFL). Like naturally occurring I1-FFL motifs in gene networks, this engineered microbial consortium acts as a pulse generator of gene expression. We demonstrated that the amplitude of the pulse can be easily tuned by adjusting the relative population fractions of the strains. We created a mathematical model for the temporal dynamics of the microbial consortium and, using this model, identified population fractions that produced desired pulse characteristics. Our work demonstrates that intercellular gene circuits can be effectively tuned simply by adjusting the starting fractions of each strain type.

## Introduction

One major goal of synthetic biology is the creation of precisely tunable gene circuits (*1*). The development of new synthetic biology tools has led to a diverse toolbox for practitioners to finely manipulate biological systems (*2*). One method for increasing the complexity of synthetic systems is to coordinate behavior across multiple interacting microbial populations (*3, 4*). Such microbial consortia exhibit several desirable properties, including division of labor, spatiotemporal organization, and emergent behavior (*5*–*7*). Division of labor is especially useful for metabolic engineering tasks as individual members of the population can specialize in a specific subset of biochemical reactions, leading to greater yield of the target product (*8, 9*). Additionally, the modular nature of synthetic consortia allows for the replacement of one member of the population with another, which can change the overall function of the population (*10*). For instance, synthetic biologists have engineered small collections of different strains that, when combined in a specific manner, can perform defined logic computations (*11, 12*). Due to the modular nature of the design, a different mixture of strains is able to perform a different logic operation without the need to redesign all strains from the bottom up.

Of particular interest are microbial consortia whose functions emerge from the interaction between the constituent strains. Examples of such circuits include population–level oscillators (*13*) and logic gates (*11, 12*). However, our ability to construct novel, engineered microbial consortia has rapidly outpaced the development of methods to tune and control their emergent function. Although usual means of tuning gene circuits, such as testing libraries of ribosome binding sites and promoters, can be applied to engineered microbial consortia, population level circuits have more parts than their monoculture counterparts. The dynamics of a multi-strain circuit are the result of the coordinated interactions of its intracellular and extracellular components. This makes both coarse systematic adjustments, and fine-tuning of desired circuit behavior more labor intensive.

Here we developed an incoherent type-1 feedforward loop (I1-FFL) that spans a population of three strains of engineered *E. coli*, and demonstrated that its behavior can be tuned by changing initial population fractions. We selected this circuit due to its interesting dynamic behavior as a pulse generator and its ubiquity in natural gene circuits (*14, 15*). We first demonstrated that pulsing behavior is observed in response to inducer only when all three strains are present using time-lapse fluorescence microscopy and bulk culture experiments. We next developed a mathematical model and fit model parameters to experimental data using Bayesian inference. Using this mathematical model, we were able to better understand the dynamic behavior of the I1-FFL circuit at different relative initial population fractions of the member strains. We found that even though the model can identify which population fractions will result in desired pulse characteristics, experimental realization was largely limited by the accuracy of strain fraction initiation. Our results demonstrate that multistrain dynamic circuit behavior can be precisely controlled by changing relative population fractions.

## Results and Discussion

### Implementation of a Multicellular Feedforward Loop

An incoherent type-1 feedforward loop (I1-FFL) consists of three nodes: X, Y, and Z (*14*). When induced X activates both Y and Z, Y represses Z after its signal accumulates above a certain threshold. This way X directly activates Z and indirectly represses it. The different dynamics of activation and repression lead to a pulse in the concentration of Z (*16*).

We constructed a multi-strain I1-FFL by engineering three strains of *E. coli* that communicate with each other using homoserine lactones (HSLs) (see Fig. 1*A*). Strain X is induced by isopropyl *β*-d-1-thiogalactopyranoside (IPTG) and produces the HSL synthase CinI, which synthesizes 3-OHC14-HSL (C14-HSL) (*17*) and mCherry2 (*18*) as a reporter (Fig. 1*B*). Strain Y is induced by C14-HSL and produces the HSL synthase RhlI, which synthesizes C4-HSL (*19*) and sfCFP. Strain Z produces sfYFP in response to C14-HSL and the RbsR-L chimeric repressor in response to C4-HSL. RbsR-L is an engineered chimeric transcription factor with a ribose responsive ligand binding domain and a DNA binding domain that targets the *lacO* operator sites of the promoter driving sfYFP (*20*). We tagged all of the fluorescent proteins, both HSL synthases, and the RbsR-L with the ssrA protein degradation tag to speed up dynamics (*21*). The complete I1-FFL is formed when the three strains are cultured together. It is activated by IPTG (Fig. 1*A*); C14-HSL acts as the activating signal that targets the second and third node; and C4-HSL acts as the indirect repression signal from the second to third node.

**Figure 1:**
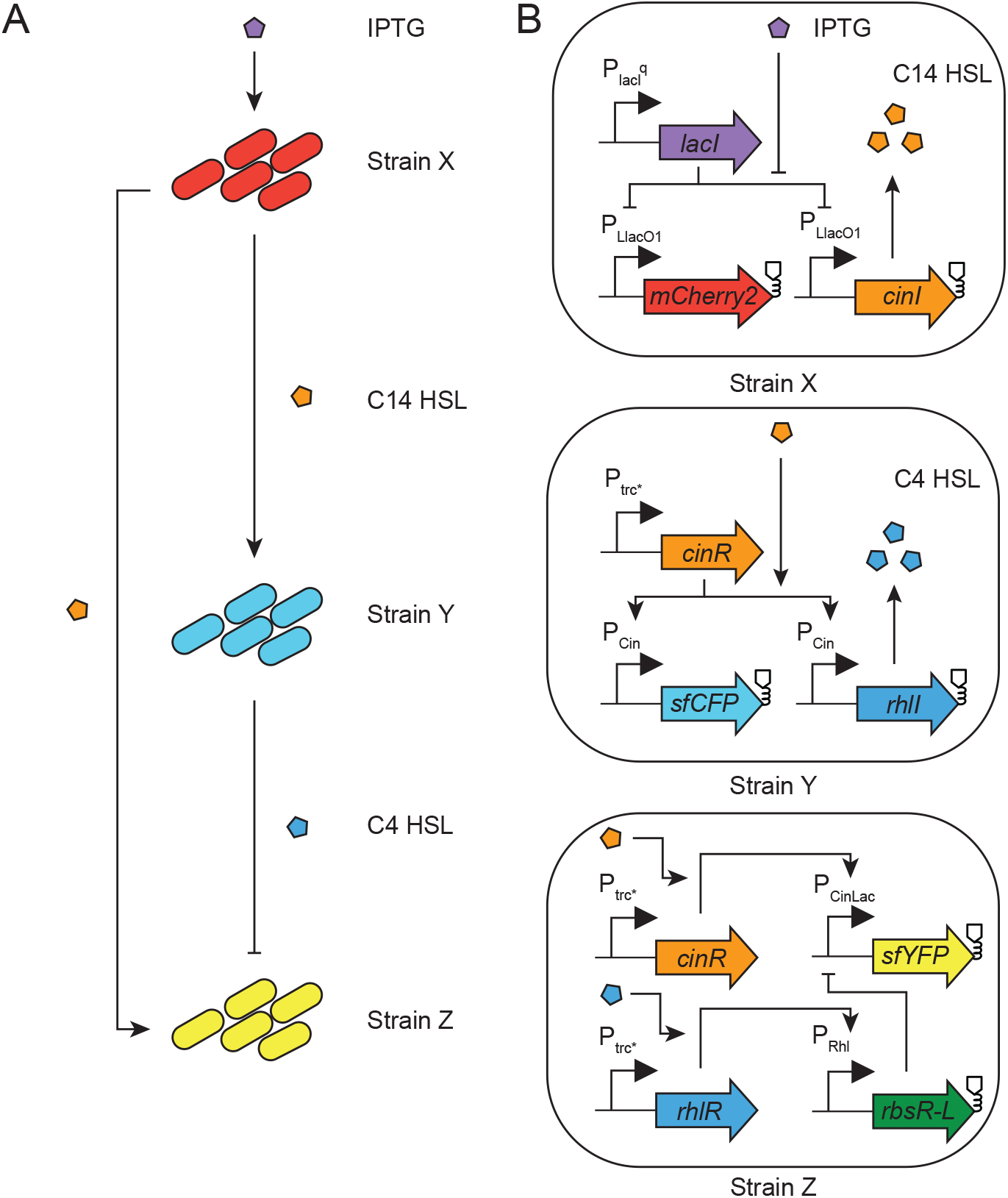
A multi-strain type I incoherent feedforward loop. (*A*) A type I incoherent feedforward loop is formed when strains X, Y, and Z are mixed together in co-culture. Orthogonal HSLs serve as signal carriers between populations. (*B*) Gene circuit schematics for strains X, Y, and Z. Proteins expressed with an *ssrA* degradation tag are labeled with the protein instability glyph.

To observe the dynamics of the consortium, we grew cells in a plate reader while measuring the fluorescence of all three strains and the optical density of the overall population over time. After the induction with IPTG we observed a rapid increase in the fluorescence signal of all three strains (Fig 2*B,C,D*). The fluorescence in strain X started to increase after induction with IPTG. This increase terminated when population reached stationary phase (Appendix Fig. S10). After this point the fluorescence signal began to decay due to the halting of transcription from σ^70^ promoters in cells that reached stationary phase (*22*). Fluorescence in strain Y showed a similar behavior to that of strain X (Fig. 2*C*). Unlike strains X and Y, strain Z generated a pulse; the corresponding fluorescence started to decay during exponential growth and by the onset of stationary phase the signal had returned close to background levels (Fig. 2*D*).

**Figure 2:**
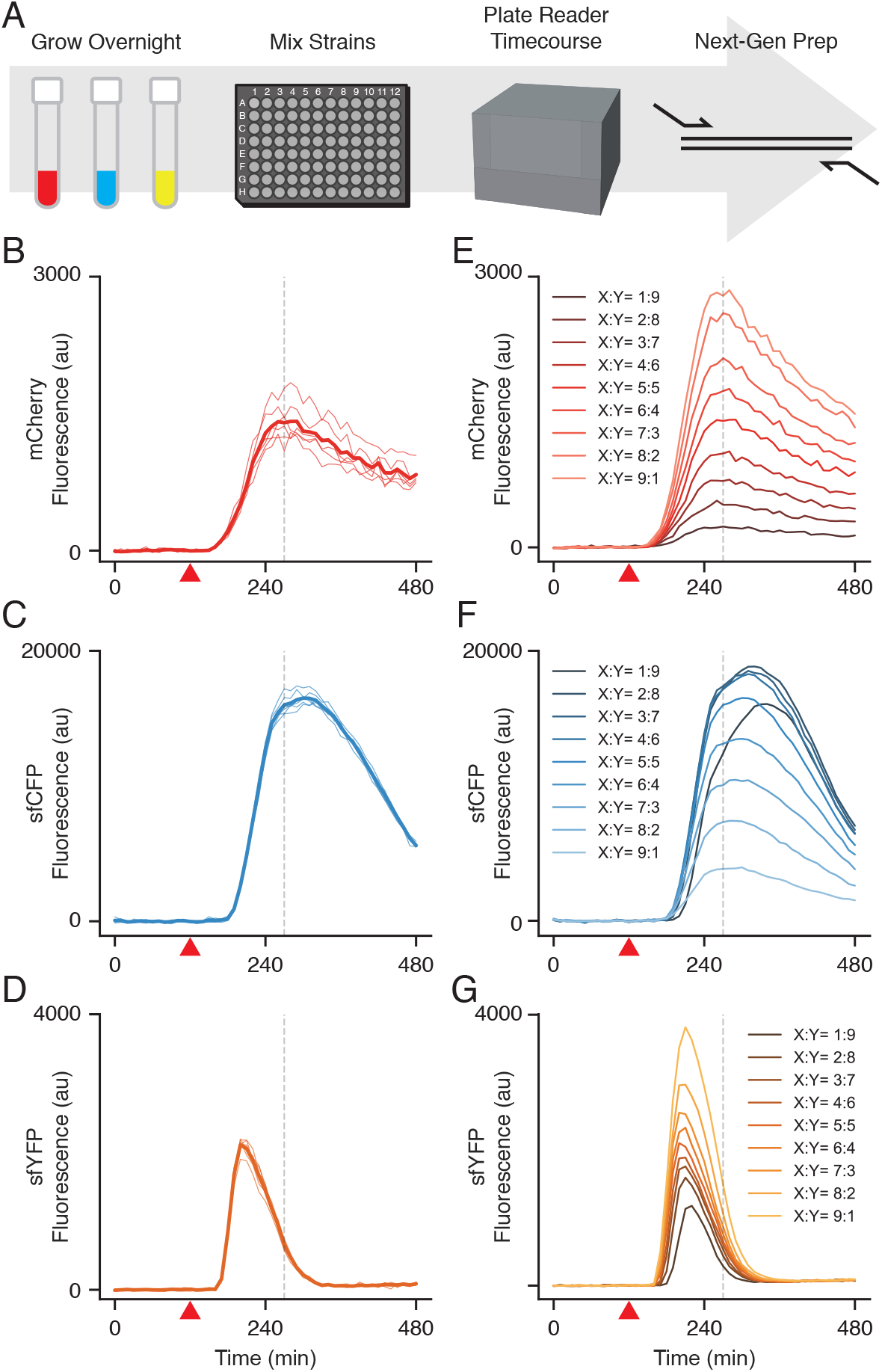
Procedure for and typical results of bulk culture experiments. (*A*) The consortial I1-FFL circuit is mixed at different relative strain fractions, measured using plate reader, and measured for strain fractions using next-generation sequencing (NGS). (*B,C,D*) Fluorescence of each strain in the I1-FFL circuit at even strain fractions during one experiment with 6 technical replicates. Thick lines represent the mean of the 6 replicates. Red triangle indicates the time at which IPTG was added (120 minutes). (*E,F,G*) Mean fluorescence activity of the I1-FFL circuit at different strain fractions. The relative fraction of strain Z was fixed to 33% and the remaining 66% of the population was divided between strain X and strain Y as indicated. Red triangle indicates the time at which IPTG was added (120 minutes). For additional replicates see Appendix Fig. S4.

To verify that the observed pulse of the Z strain was not due to artifacts introduced by the reduced protein expression in bacterial populations approaching stationary phase, we probed the three strain consortium using microfluidics. The microfluidic device used in this experiment has a large, 2 mm long trapping area, which increased the likelihood that all three strains were present and co-localized during the experiment (Appendix Fig. S1) Fluctuations in strain proportions can lead to incidental extinction in smaller trapping regions. Larger trapping areas decrease variability in population fractions, and thus reduce the risk of such extinctions (*23*). Upon induction with 2 mM of IPTG, all three strains responded by a rapid increase in fluorescence signal. However, 48 minutes after induction, the fluorescence signal of strain Z began to decrease, in contrast with the fluorescence of strains X and Y, which continued to increase (Appendix Fig. S2). This result shows that the consortial I1-FFL circuit generates a pulsing behavior of the Z strain in the microfluidic setting, confirming the results obtained with the bulk culture experiments.

To additionally verify that the observed behavior of the consortial I1-FFL circuit was indeed generated by the correct circuit topology we built alternate circuits that are not expected to pulse. As expected, these consortia do not allow for the generation of a pulse in the Z strain (Appendix Fig.3). Taken together, these results demonstrate that our consortial I1-FFL circuit has a robust behavior, able to generate pulses in the Z strain in both microfluidics and bulk culture assays. Moreover, the observed pulse is specific to the distributed genetic circuit topology.

We hypothesized that the relative fraction of each strain in the microbial consortium influences the overall behavior of the circuit because it affects the amount of HSL molecules in the culture. To prove this hypothesis, we performed a bulk culture experiment in which we changed the ratio of the three strains of the consortium at the time of innoculation; We kept the fraction of strain Z fixed to 33% of the population and assigned different fractions of the remaining 67% of the population to strains X and Y. We chose the bulk culture assay because, unlike microfluidics, it offers better control over strain seeding and allows a uniform distribution of HSL molecules in the culture Fig. 2*E,F,G* show the average time course of the fluorescence signal of the strains X, Y, and Z, respectively, computed from 6 technical replicates (variability between technical replicates is similar to that in Fig. 2*B,C,D*). The peak mCherry signal increased as the fraction of strain X was increased. The sfCFP signal also increased as the fraction of strain Y was increased up to an X:Y ratio of 5:5, upon which the sfCFP amplitude reached a maximum. At X:Y ratio 1:9, the magnitude of the peak in Y fluorescence decreased despite having a larger population fraction because the smaller number of strain X cells resulted in slower accumulation of the C14-HSL signal that drives protein synthesis in strain Y. This demonstrates that the output of strain Y is dependent on the strain fraction of both strain X and strain Y. At high X:Y ratio a greater activating signal from strain X combined with a reduced repressive signal from strain Y, resulting in a higher pulse amplitude in strain Z. These results demonstrate that the system responds to changes in strain fraction.

To determine if the fractions of each strain in the culture matched our target fractions, we performed next-generation sequencing (NGS) on samples before (t = 0 min) and after (t = 480 min) the time course experiment. A unique 25 bp barcode was integrated into the genomes of strain X, Y, and Z at a single locus.The barcodes were amplified off the genome using PCR, and sequenced on an HiSeq platform (Appendix Fig. S5). The fraction was then calculated by counting the occurrence of each barcode and dividing by the total barcodes counted. We found that the measured fractions of cells did not fluctuate significantly over the course of the experiment (Appendix Fig. S8). However, we found that the measured fractions at the start of the timecourse reported by the NGS experiments differed from the target fractions, specifically, the fraction of strain X was consistently lower (Appendix Fig. S6).

### Mathematical Modeling of Multi-Strain I1-FFL Dynamics

Our next goal was to capture the dynamics of the multi-strain incoherent feedforward loop using a mathematical model. To do so, we developed a low-dimensional system that describes the essential interactions in the circuit.

Our model describes the bulk average fluorescence and quorum sensing signals, but not the cell to cell variability across the consortium. The number of cells in our experiments is large, so that variability between individual cells is averaged out in the signal. Hence, a mean-field description is appropriate. We also neglected spatial effects, such as the diffusion of the signaling molecules, because experimental wells were shaken to keep the contents well-mixed. The resulting mathematical model has the following form:

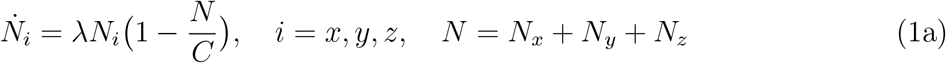

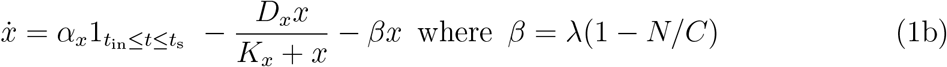

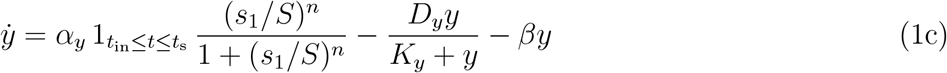

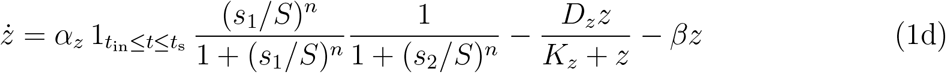

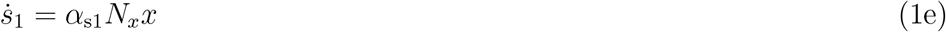

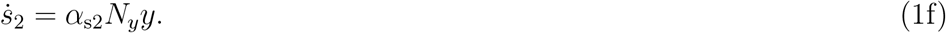

Equation 1a describes the cell growth dynamics with *N*_*x*_, *N*_*y*_, and *N*_*z*_ as the population sizes of strains X, Y, and Z, respectively, and *N* as the total number of cells. The parameters *λ* and *C* represent, respectively, the cell growth rate, and the carrying capacity of a well.

Equations 1b-1d describe the dynamics of the proteins in each strain, where *x, y*, and *z* are the intracellular protein concentrations in strains X, Y, and Z, respectively. We assume that *x, y*, and *z* also represent the concentrations of the corresponding fluorescent proteins in each strain, as the fluorescent protein is co-expressed with the corresponding synthase. To model the dynamics of each protein, we used a synthesis term, an enzymatic degradation term, and a dilution term. Protein synthesis rate in strain X is modeled by an indicator function, 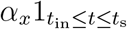, representing induced gene expression starting after induction at time *t*_in_, and ending upon reaching the cell saturation phase at time *t*_s_. Although protein synthesis dynamics may be complex during stationary phase, we assumed that synthesis ceases completely for simplicity. This simplification is reasonable for the purposes of our model because the pulse in strain Z occurs during the exponential cell growth phase. Protein synthesis in strains Y and Z are modeled similarly, but are multiplied by regulatory functions representing the activation and inhibition by quorum sensing signals. Finally, Equations 1e-1f describe the synthesis of HSLs by the corresponding synthases, where *s*_1_ and *s*_2_ represent the concentrations of C14-HSL and C4-HSL, respectively. Further details about the model, including an explanation of all parameters, see the section “Construction of the mathematical model” in Methods.

As noted earlier, NGS indicates that the fractions of strains X, Y, and Z do not fluctuate over the course of the experiment (Appendix Fig. S8), but deviated from our target strain fractions (Appendix Fig. S6). We therefore used the fractions measured using NGS at the onset of the experiments in our mathematical model. We write *N*_*x*_ = *r*_*x*_*N, N*_*y*_ = *r*_*y*_*N*, and *N*_*z*_ = *r*_*z*_*N*, where *r*_*x*_, *r*_*y*_, and *r*_*z*_ are the fractions of strains X, Y, and Z, respectively. We can tune the pulse generated in strain Z by varying these strain fractions as shown in Fig. 2.

### Model Fitting and Validation

We next fit Eq. (1) to experimental data, and used the resulting model to predict the behavior of the circuit across different strain fractions. We then validated these predictions with further experiments. First, we inferred the unknown model parameters using the measured time course OD and fluorescence data from 3 representative experiments performed with different strain fractions: (*r*_*x*_ = 0.2, *r*_*y*_ = 0.47), (*r*_*x*_ = 0.47, *r*_*y*_ = 0.2), and (*r*_*x*_ = 0.2, *r*_*y*_ = 0.2). We then used Bayesian inference (*24*) to fit the model to data. For more details about our Bayesian approach, see the section “Bayesian Parameter Inference” in Methods. Fitting to data from experiments at a single strain fraction, rather than at three different fractions, provided reasonable, but less accurate predictions (See Appendix data Fig. S9.)

Next, we designed two additional experiments to assess the predictive power of our model. In particular, we used the model to predict pulse intensities in strain Z as a function of the strain fractions *r*_*x*_ and *r*_*y*_ (Fig. 3*D*). In the first experiment, we set the fraction of strain X at 0.2 and chose fractions of strain Y predicted to produce regularly spaced peak intensities (black dots in Fig. 3*D* show the fractions we used in the experiments). Fig. 3*A,B,C* show the experimentally measured fluorescence in strains X, Y, and Z, respectively. Because the fraction of strain X remained fixed across conditions, its dynamics remained unchanged, as expected. Fluorescence of strain Y increased in strength with an increase in its fraction, *r*_*y*_. For strain Z, we observed approximately regularly spaced peak intensities. Experimentally observed fluorescence intensities followed the predictions of the model closely. For instance, the experimentally obtained results match the predicted peaks (Fig. 3*E*). The model was also able to predict the trends in the timing of the peak observed experimentally: Compare the experimental time course data in Fig. 3*C* with model simulation results in Fig. 3*F*.

**Figure 3:**
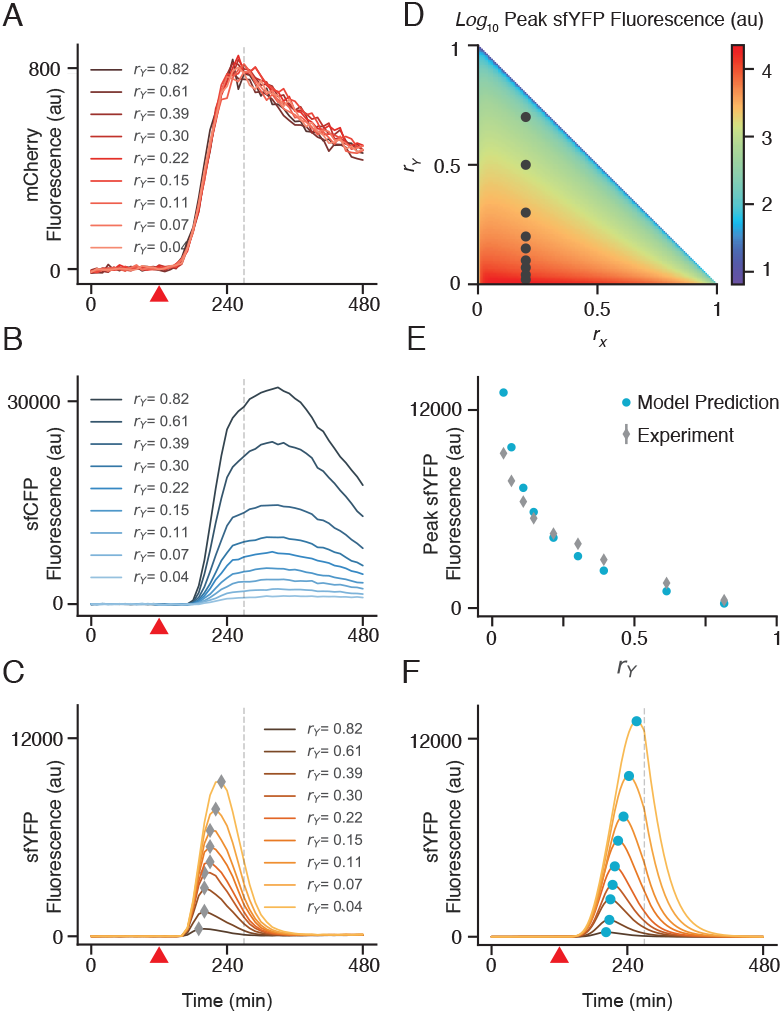
Experimental validation of the model. Strain fractions were selected by fixing the population fraction of X, and varying the fractions of Y and Z. Each condition is distinguished by the population fraction of Y. (*A,B,C*) Fluorescence of each strain of the I1-FFL system at 9 different population fractions. Peak heights are annotated in the strain Z. Red triangle indicates the time at which IPTG was added (120 minutes). (*D*) Simplex of peak height as a function of relative fraction of X and Y. Target fractions selected for the experiment are displayed as black dots. (*E*) Mathematical model predicts the peak height as a function of strain fraction measured at experiment onset using NGS. Error bars on experimental data represent standard deviation across 6 technical replicates performed on the same day. Error bars are smaller than the markers so they are not visible. (*F*) Model simulation predicts amplitude of the pulse in strain Z as well as temporal behavior as a function of strain fraction. Red triangle indicates the time at which IPTG was added (120 minutes). For additional replicates see Appendix Fig. S7.

The close agreement of experimental data and model predictions suggests that the model captures the main mechanisms that drive the circuit’s dynamics. However, model predictions deteriorate at smaller fractions of strain Y (compare experimental and simulation results for *r*_*Y*_ *<* 0.15 in Fig. 3*C,F*). This disagreement may be due to the limited ability of the model to capture the behavior of the system at extreme strain fractions. When one of the populations is much smaller than the others, the assumption that they are well-mixed may not hold, and stochastic effects may become significant. Our deterministic model does not capture these effects. Further, it may also be that either pipetting such small fractions or measuring those fractions *a posteriori* (or both) is not accurate enough to provide the model with the correct initial condition. Moreover, the posterior distributions over the model parameters also revealed that the growth rates were strongly constrained by the data, while the enzymatic degradation rates were constrained more weakly (See Appendix Fig. S11).

### Model Guided Pulse Generation

Next we asked whether we can generate a pulse in strain Z fluorescence with desired amplitude by using our model as a guide. We selected a wide range of strain fractions that were all predicted to generate peak fluorescence intensity in the strain Z of approximately 1700-1750 a.u. (black circles in Fig. 4*A*). To account for the error when setting initial strain fractions, we fit a linear model to the next-gen sequencing results and used the model to calculate a correction to account for the error (Appendix Fig. S12). This resulted in measured strain fractions that are significantly closer to the targeted strain fractions than previous experiments, shown here are strain fractions of the experiment shown in (Fig. 2*E*) without the correction applied (Fig. 4*B*). The dynamics of strain X again depended on the fraction of strain X, with stronger responses at higher fractions. We observed the same trend with strain Y, except at very small fractions of strain X, which caused a delay in the rise of the fluorescence in strain Y due to additional time needed for the generation of a sufficiently strong signal.

**Figure 4:**
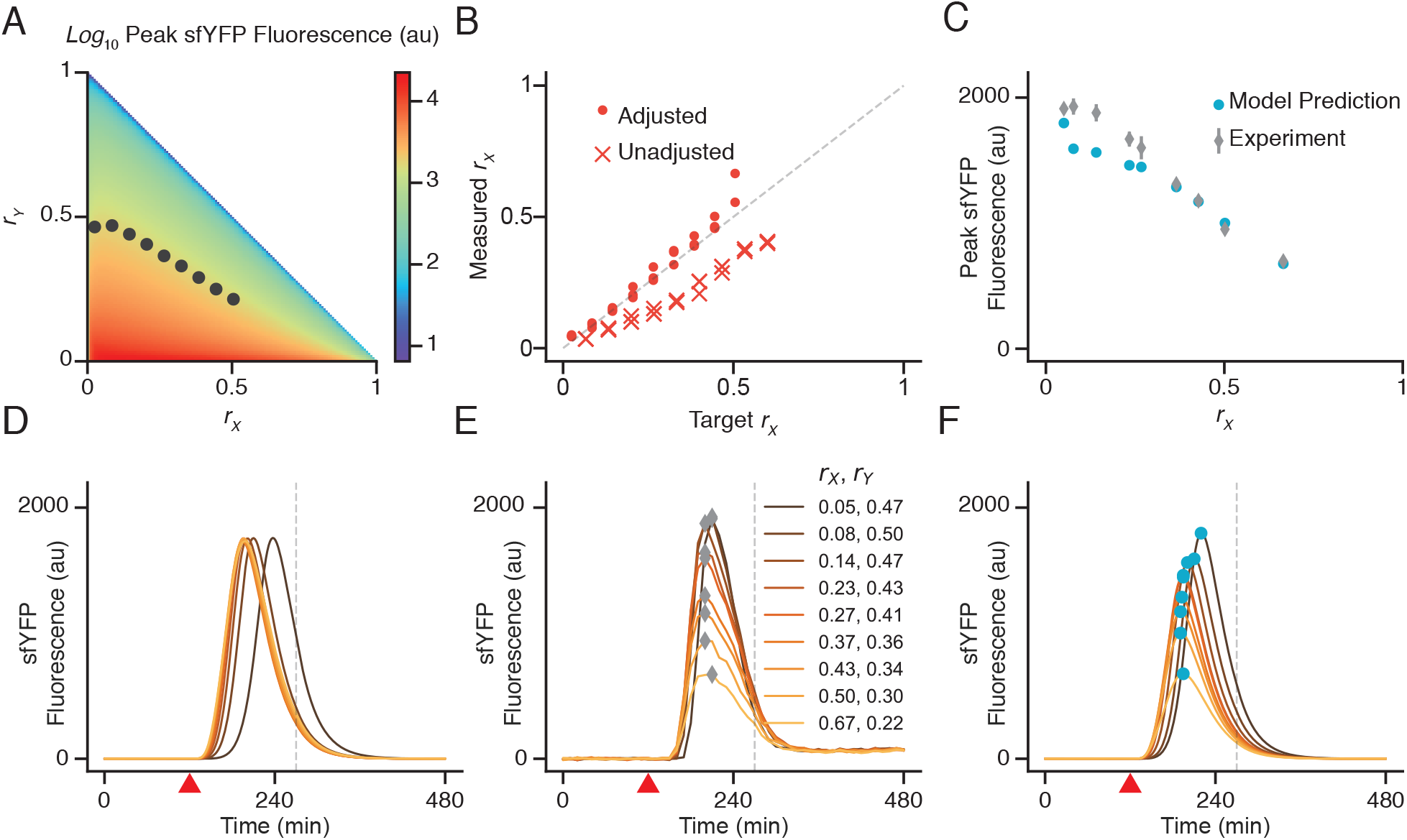
Generating pulses of constant height. (*A*) We selected strain fractions from the simplex that were predicted to have the same peak height and performed a bulk culture experiment using these strain fractions. To accurately pipette the required strain fractions, we performed a correction to account for systematic error in strain fraction observed in previous experiments. (*B*) Target versus measured strain fractions of experiments with the correction applied and without the correction applied for strain X (*C*) Model prediction of peak height based off of measured strain fraction compared to experimental data. Error bars represent three standard deviations of 6 technical replicates.(*D,E,F*) Fluorescence of strain Z. (*D*) Model simulation based off of target fraction, (*E*) Experimental data with peaks labeled using grey diamonds, (*F*) Model simulation based off measured fraction with peaks labeled using blue circles. Red triangle indicates the time at which IPTG was added (120 minutes).

At small values of the fraction of strain X, strain Z peak intensities were relatively close to the 1700 a.u.value predicted by the model using target strain fractions. However, at larger fractions, the peak fluorescence declined. Although after applying the correction the measured strain fraction was closer to the target, we conjectured that the decline in peak fluorescence was due to the measured strain fractions differing from the target fraction. We confirmed this by generating model predictions using the measured strain fractions as input (Fig. 4*C*). Moreover, the peak of the pulse is more sensitive to changes in strain fractions at larger values of the fractions of strain X (note that the peak isoclines converge in the right corner of the simplex).Therefore a small change in strain fractions can considerably change the peak intensity. By using the measured strain fraction values obtained by NGS, our model predicted the experimentally observed decline in peak fluorescence (Fig. 4*C*), however these predictions deviated from the predictions made using target strain fractions (Fig. 4*D*). Our model was also able to predict the finer details of fluorescence dynamics, as well as the shifts in the timing of the peak (see Fig. 4*E, F*). We thus concluded that the model is able to predict the dynamics of the strains, given that the strain fraction is known.However, to control these dynamics, one needs to accurately set the initial strain fractions, which can be difficult especially if one of those fractions is small.

## Discussion

Here we presented a highly tunable multi-strain synthetic gene circuit along with a simple model that accurately predicts the circuit’s dynamics.We found that tunability came at the price of robustness to perturbations in strain fraction: Strain fractions could deviate from their target values by as much as 20% of the total, causing pulse amplitudes to miss their predicted values. Furthermore, due to the sensitivity of the system, even small deviations in strain fractions leads to significant differences in experimentally measured peak heights. The model revealed when peak height as a function of strain fraction was flat. At these strain fractions the system was robust to fluctuations in strain fraction, but exhibited a reduced range of peak amplitudes. The model also correctly predicted that in other regions of the fraction space, amplitude can be significantly affected by fraction deviations as low as 1%.

This trade-off between robustness and tunability is a ubiquitous problem for the design of synthetic circuits. Often, nature’s solution to the problem is the addition of negative and positive feedback (*25*). Synthetic circuits that are designed to be both robust and tunable take advantage of feedback control to reject perturbations while remaining responsive to changes in their inputs (*26, 27*). This suggests the next step in circuit design: The inclusion of feedback control to self-regulate population fractions which will then, in turn, control circuit behavior. For instance, one could engineer a cell lysis module such that when one of the strain fractions begins to deviate from its target value, the module initiates cell suicide in that strain, rebalancing the strain fractions (*28*). This suggests that one could introduce genetic modules to encode fixed values for population fraction, enabling our circuit to pulse at fixed pulse amplitude given any initial seeding fraction.

One advantage of our tuning approach is the simplicity and accuracy of the model, despite inaccuracies in strain seeding. Our model only requires fitting to just a few experimental runs and can accurately predict the circuit’s dynamics over most of the strain fraction space. A potential drawback of this approach is that the method fails when strain fractions change significantly over the course of the experiment. This could be caused by a difference in growth rate or an inability to accurately set the initial strain fractions, as we have demonstrated. Another advantage is that using relative strain fractions to tune circuit behavior is relatively easy compared to more traditional tuning approaches, such as making ribosome binding site libraries followed by rounds of high throughput screening. In our experiments, the three strains were cloned once with no further iterations, yet produced a vast range of predictable dynamic behaviors. By shifting the “atomic unit” of tuning to the population fraction of the strain, behavior need not be tuned by altering DNA, but instead by simply mixing appropriate amounts of cell culture.

## Methods

### Fabrication of Microfluidic Device

The microfluidic device was created using the protocols of Ferry et al. (*29*). The device is a modified version of the 2000 μm open device described in Alnahhas et al. (*23*) with a Dial-a-Wave junction and chaotic mixer channels for media mixing (*30*). See Appendix Fig. S1 for details.

### Strains and Plasmids

All strains used in this study were derived from CY27 (BW25113 Δ*sdiA* Δ*lacI* Δ*araC rhlR::cinR*) (*13*). DZ02 was created by inserting *lacI* driven by the *PlacIq* promoter at the safe site 9 locus (*31*) of CY27 using lambda red recombineering (*32*). To uniquely label strains for next-gen sequencing, we designed barcodes for strains X, Y and Z by choosing three 25 bp sequences from a collection of 25 bp DNA probes (*33*). DZ06 was created by inserting a 200 bp segment of DNA containing the barcode for strain X at safe site 3 locus of DZ02. DZ07 and DZ08 similarly were integrated with the barcodes for strain Y and Z respectively at the safe site 3 locus of CY27.

All plasmids were assembled using Golden Gate Assembly (*34*). pDZ149 contains two expression cassettes, one expresses CinI and the other expresses mCherry2 (*18*), both from the P_LlacO1_ promoter. pDZ047 is an empty vector that contains ampicillin resistance cassette and the p15A origin of replication. Strain X is created by co-transforming pDZ149 with pDZ047 in strain DZ06. pDZ033 contains two expression cassettes, one expresses RhlI and the other expresses sfCFP, both from the P_CinLac_ promoter (*13*) and both proteins are tagged with ssrA. Strain Y is created by co-transforming pDZ033 with pDZ047 in strain DZ07. pDZ114 contains one expression cassette which expresses sfYFP tagged with ssrA driven by the P_CinLac_ promoter. pDZ134 contains one expression cassette which expresses RbsR-L driven by the P_RhlTet_ promoter. Strain Z was created by co-transforming pDZ114 with pDZ134 in strain DZ08. pDZ113 and pDZ115 were used in creating control strains shown in Appendix Figure S3. pDZ113 contains on expression cassette which expresses sfYFP from the P_Rhl_ promoter. Strain Z Cascade was created by co-transforming pDZ113 with pDZ047 in strain DZ08. pDZ115 contains on expression cassette which expresses sfYFP from the P_LlacO1_ promoter. Strain Z Inverter was created by co-transforming pDZ115 with pDZ134 in strain DZ08. All expression cassettes use BCD bicistronic design ribosome binding sites. See Appendix Table S1 for summary.

### Microscopy

Glycerol stocks of transformed strains for the I1-FFL were streaked onto an LB Agar (Genesee) plate containing 100 mg/L ampicillin (Fisher BP1760) and 50 mg/L kanamycin (Acros Organics BP906). The plate was then incubated in a stationary incubator overnight at 37 °C, 16-18 hours. The following day, the plate was removed from the incubator, parafilmed (Bemis), and refrigerated at 4 °C for further use. One 3 mL overnight liquid culture was prepared for each cell strain using LB media with 100 mg/L ampicillin and 50 mg/L kanamycin in plastic 14 mL culture tubes. The tubes were placed in a 37 °C incubator with orbital shaking at 250 rpm for 16-18 hours overnight. Each culture was diluted 1:1000 into 15 mL of fresh LB media with 100 mg/L ampicillin and 50 mg/L kanamycin in 25 mL shake flasks. These cultures were grown until they reached an OD600 of 0.1 as read using a UV-Transparent Disposable Cuvette (BrandTech 759150) in a NanoDrop 1000 Spectrophotometer. The cultures were then transferred to a 15 mL conical tube and incubated on ice for 5 minutes. Next, the cultures were centrifugated at 2000 x g for 10 minutes at 4 °C. The supernatant was decanted. The pellet of the first strain was then resuspended in 10 mL of LB containing 100 mg/L ampicillin and 50 mg/L kanamycin. This resuspension was then used to resuspend the second pellet and then that resuspension was then used to resuspend the third pellet. This results in an even mixture of all three strains in a final volume of 10 mL and an approximate OD600 of 0.3-0.4. Next the cells were loaded into a 10 mL syringe with luer stub attachment and Tygon tubing. Syringes for the induced media (LB containing 2 mM IPTG, 0.01% Tween 20, 100 mg/L ampicillin, 50 mg/L kanamycin, and 10 mM D-Ribose (Sigma-Aldrich R7500)), uninduced media (LB containing Sulforhodamine 101 at 200 ng/mL, 0.01% Tween 20, 100 mg/L ampicillin 50 mg/L kanamycin, and 10 mM D-Ribose) in 50 mL syringes, and water balance/waste were also created (4 water syringes total in 10 mL syringes). Syringes were then affixed to the sliding rail for fixed heights, or the linear actuator for the dynamic heights. Tubing was then plugged into the microfluidic device in the following order: Induced Media, Water Balance, Uninduced Media, Waste 1, Waste 2, and Dummy Cells (a placeholder containing water). Heights for the linear actuator for induced and uninduced states were then programmed using custom software (*29*). Next, the time course program was then input in the custom software as follows: square wave with period 12 hours, amplitude of 100%. The Dummy Cell syringe was then unplugged from the device and replaced with the cells syringe. To facilitate trapping, the tubing line was agitated by flicking. As many cells were put into the trap as was possible to increase the likelihood that neighboring cells are of different cell types. Flow was reversed by lowering the height of the cells until the flow rate in the open channel reached approximately 100 μm per second. Cells were grown in the uninduced media condition until the trap was visibility full or close to filling. At this time, the time course program is started, and image acquisition begins. Acquisition was performed on a Nikon Ti-E inverted fluorescence microscope with Perfect Focus at 37 °C. The image acquisition settings are as follows: 11 fields of view horizontal stitched together, image taken every 6 minutes, 4 channels of acquisition in the following order: 60x Phase Contrast, mCherry, sfYFP, sfCFP. Image acquisition continues until acquisition is stopped manually.

### Microplate Timecourse Experiments

Glycerol stocks of transformed strains were streaked onto an LB Agar (Genesee) plate containing 100 mg/L ampicillin (Fisher BP1760) and 50 mg/L kanamycin (Acros Organics BP906). The plate was then incubated in a stationary incubator overnight at 37 °C, 16-18 hours. The following day, the plate was removed from the incubator, parafilmed (Bemis), and refrigerated at 4 °C for further use. One 3 mL overnight liquid culture was prepared for each cell strain using M9CA media (M9 media with 0.2% Cas amino acid, 0.4% glucose, 100 mg/L ampicillin and 50 mg/L kanamycin) in plastic 14 mL culture tubes. The tubes were placed in a 37 °C incubator with orbital shaking at 250 rpm for 16 hours overnight. The following day, the Tecan Spark (Tecan Trading AG) was preheated to 37 °C. While the Tecan Spark warms, contents of the 96 well plate was pipetted. The overnight cultures were diluted 1:10 and measured OD600 in a UV-Transparent Disposable Cuvette (BrandTech 759150) in a NanoDrop 1000 Spectrophotometer, the OD was recorded and then diluted to a final OD600 of 0.005. Each strain was mixed to the strain fraction as needed by the experiment by volume and pipetted into the wells of a Corning 96-well Flat Clear Bottom Black Polystyrene TC-treated Microplate (Corning 3904). For the same peak height experiment, calculations for the volume to pipette were made based off of next-gen sequencing data. For more information see section. A small amount of the pipetted mixture was frozen at -20 °C. The plate was sealed with a Breathe-Easy (Diversified Biotech) sealing membrane, ensuring that there were no gaps between the seal and the wells (See discussion on the use of the plate seal). The following settings for the Tecan Spark were used: temperature set to 37 °C, kinetic loop for 8 hours, orbital shaking at 510 rpm, acquire events every 10 minutes. The following events were collected with the following settings: OD (Wavelength: 600 nm), mCherry2 (Ex: 580 nm, Em: 611 nm, bandwidth 5 nm, gain: 190), sfCFP (Ex: 455 nm, Em: 472 nm, bandwidth 5 nm, gain: 190), sfYFP (Ex: 517 nm, Em: 537 nm, bandwidth 5 nm, gain: 170, number of flashes: 30). Data were acquired for 2 hours and paused button after cycle 12 completes reading but before cycle 13 initiates. The plate seal was removed. IPTG (RPI 367-93-1) was added to a final concentration of 2 mM, we use 2 μL of 200 mM IPTG. The plate was sealed again a Diversified Biotech Breathe-Easy sealing membrane, ensuring that there are no gaps. The plate was returned to the Tecan Spark and the acquisition loop was allowed to continue. When the acquisition was completed the plate was removed from the Tecan Spark and the plate seal was removed from the plate. An aluminium plate seal (Diversified Biotech) was applied and the plate was frozen at -20 °C.

### Note About Plate Seal Usage

To reduce the effect of evaporation over long time frames we used a semipermeable plate seal, specifically, the Breathe-Easy (Diversified Biotech) sealing membrane. We found that the use of this plate seal exacerbated the micro-aerobic environment of the 96 well plate, causing the cells to enter stationary phase due to oxygen starvation. Typically the cells to enter stationary phase at around 270 minutes. It is also likely that low oxygen concentrations led to reduced fluorescent protein maturation and lower fluorescence signal.

### Next-gen Sequencing Experiment

Strains DZ06, DZ07 and DZ08 were modified with a unique 25bp barcode for identification using amplicon sequencing. This 25bp barcode, along with surrounding regions from the knock in cassette are located on the genome at the SS3 locus (*35*). Primers were designed to bind to regions flanking the unique barcodes such all three barcodes could be amplified using one primer pair in a single PCR reaction. Primers contain 3 regions. From 3’ to 5’: a homology region that anneals to the genome, a 5 bp barcode that identifies the primer, and a partial Illumina adapter sequence that is required for the Amplicon-EZ service. Since each microplate experiment for varying strain fractions uses 54 wells (6 technical replicates of 9 different conditions), we designed n=6 5bp barcodes on the overhang of the forward primer, and m=9 5bp barcodes on the overhang of the reverse primer. Each primer pair corresponds to a specific well in the microplate experiment. The end of forward and reverse overhangs contains the partial Illumina adapter sequences required by Genewiz’s Amplicon-EZ service. For each experiment, samples were frozen at -20 °C at time t=0 minutes and t=480 minutes. For samples at t=0, 20 μL of liquid culture mixed to specified fractions at 0.05 OD were frozen (See Microplate Experiments). For samples after the end of the experiment, the 96 well microtiter plate was sealed with film and frozen (See Microplate Experiments). For PCR, the samples were thawed on ice and 1 μL of the culture was added to a 20 μL PCR reaction using NEB Q5 polymerase and GC Enhancer. 9 PCR reactions were required for time points before the start of the experiment, while 54 were required for time points after the end of each experiment due to the 6 technical replicates. The initial denaturing step of the PCR was set to 10 minutes to lyse the cells and reveal the genomic DNA. Next, the cycles are as follows: 98 °C for 10 seconds, 65 °C for 20 seconds, 72 °for 10 seconds. This was repeated for 25 cycles to prevent post exponential amplification bias. Following the 25 cycles: a final elongation step at 72 °C for 5 minutes and an infinite hold at 12 °C. PCR reactions from the same time point were then pooled together and purified using a spin column DNA clean and concentrate kit (Zymo). To prevent oversaturation of the spin column, a maximum of 6 reactions were pooled together. Purified amplicons were then measured using Nanodrop and Qubit, normalized to 20 ng/μL and sent to Genewiz for Amplicon-EZ sequencing service (Illumina HiSeq 2×150). This service typically returns 40,000 sequences however in some cases fewer than 40,000 sequences were returned. FASTQ sequencing files were analyzed using a Python script (see Github for source code).

https://github.com/biodesign-lab/AmpliconRatios

In summary, first, exact matches were used to find the 5 bp barcodes that identify the well the sample came from as certain mismatches can alias to another 5 bp barcode. Next, an alignment is performed with the known sequence of each of the 3 barcodes. This process is repeated for every entry in the FASTQ file. Typically, around 7% of sequences are rejected.

### Plating Assay

The plating assay was performed according to the methods described in Molinari et al. (*36*).

### Calculation of Pipetted Fractions

We found that strain X had consistently lower strain fraction (measured by NGS at experiment onset) relative to target fractions (Appendix Fig. S6). This is consistent with the results of the plating assay because strain X was found to have a lower CFU/OD600 (Appendix Fig. S13). Therefore, setting strain fractions by normalizing by OD600 may have caused less of strain X to be pipetted into the plate than the target amount. For the experiment that keeps the peak height the same we used the next gen sequencing data to generate a linear transformation of the volume to pipette. For the results of this linear transformation, see Appendix Fig. S12.

### Construction of the Mathematical Model

Below, we provide the mathematical model of the consortial pulse generator and a description of different terms and parameters in the model. The dynamics of the consortial pulse generator are described by the following set of ordinary differential equations

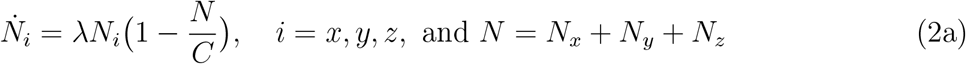

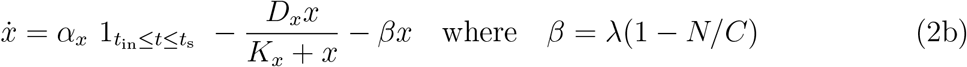

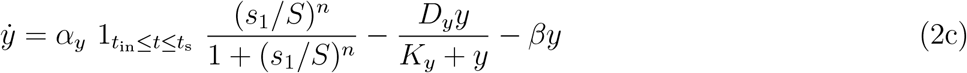

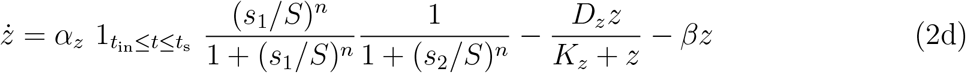

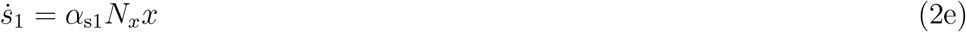

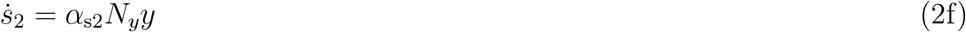

Equation (2a) describes the cell growth dynamics where *N*_*x*_, *N*_*y*_, and *N*_*z*_ represent the cell populations of strains X, Y, and Z, respectively. The parameter *λ* is the cell growth rate coefficient and *C* is the carrying capacity of the container the cells grow in. Note that *N* = *N*_*x*_ + *N*_*y*_ + *N*_*z*_ is the total cell population in the liquid culture and the instantaneous cell growth rate is *β* = *λ*(1 *− N/C*) that satisfies the equation 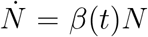. In particular, note that when the total cell count reaches the proximity of the carrying capacity of the container *C*, the instantaneous cell growth rate approaches zero. Equation (2a) is a logistic growth model that approximates the dynamics of growth in microplate wells. The cells first grow in an exponential phase, but ultimately reach a stationary phase due to nutrient depletion and toxic waste buildup. In the model, cells grow with a rate of order *β ≈ λ* when *N* is small, but when *N* gets close to *C*, the growth rate approaches zero.

We assume all cells grow with the same effective rate *β*. This implies that the relative fraction between strains will remain constant throughout the experiment. In other words, the fractions *r*_*x*_, *r*_*y*_, and *r*_*z*_ of strains X, Y, and Z, respectively, do not change in time. We have tested this assumption by measuring the strains fractions using the NGS technique described in Appendix Fig. S5 at the beginning and the end of the experiments. We saw that the relative fractions between strains remained approximately constant (Appendix Fig. S8), and we thus assumed that the strain fractions remained fixed during the experiment.

Equation (2b) describes the dynamics of the protein synthesis and degradation in strain X. In particular, *x* denotes the average cellular concentration of both the synthase CinI and the fluorescent protein mCherry2. The assumption that the synthase and the fluorescent protein concentrations are equal is justified as these two proteins have the same promoters and are degraded with the same degradation tags. In reality, at the single cell level, different sources of noise, such as variability in transcription, translation, and degradation, result in difference in the concentrations of the synthase and fluorescent proteins. Since here we model average concentrations across cells, we effectively assumed that the synthase and fluorescent protein concentrations are approximately equal.

The protein synthesis term in strain X is 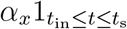 which models induced gene expression starting at time *t*_in_ (the induction time) and shuts off at time *t*_s_ (the start of the stationary phase). The assumption that the gene expression shuts off as the stationary phase of the cell growth begins was made for simplicity. In particular, the gene expression dynamics in the stationary phase may be complicated as the cells run out of the nutrients and start using secondary carbon sources. On the other hand, our primary focus is the dynamics and tuning of the pulse generated in strain Z fluorescence levels. Given that this pulse occurs (for the most part) in the exponential phase of the cell growth (see Fig. 2**g** for instance), modeling the dynamics of the stationary phase is unnecessary for our purpose.

The parameter *α*_*x*_ is the protein synthesis rate in strain X. The term *D*_*x*_*x/*(*K*_*x*_ + *x*) is a degradation term modeling the enzymatic degradation of protein *x* using ClpXp. This term can be obtained by assuming a Michaelis-Menten mechanism for the enzymatic degradation of the protein (*37*). The parameter *D*_*x*_ is the degradation rate coefficient and *K*_*x*_ is a Michaelis-Menten constant. The term *βx* accounts for dilution due to cell growth at rate *β*. We assume that the total volume occupied by cells is proportional to the total number of cells.

Equation (2c) describes the dynamics of protein *y* in strain Y (*i*.*e*. the synthase rhlI and the fluorescent protein sfCFP). The dynamics are similar to those of protein *x*, except that the protein *y* synthesis rate is multiplied to a regulatory (Hill) function that captures the activation of the protein synthesis in strain Y by the quorum sensing molecules emitted from strain X, *s*_1_. The parameter *S* is the amount of quorum sensing signal needed to make the protein synthesis rate half of the potential maximum. The exponent *n* reflects the sensitivity of the protein synthesis rate to the quorum sensing signal.

Equation (2d) describes the dynamics of protein *z* in strain Z (*i*.*e*. the fluorescent protein sfYFP). Since the protein synthesis rate in strain Z is activated by the signaling molecule from strain X (*s*_1_) and inhibited by the signaling molecule from strain Y (*s*_2_), the protein synthesis rate in strain Z is multiplied by two regulatory Hill functions, one representing the activation by *s*_1_ and the other representing the inhibition by *s*_2_. The exponent *n* is the Hill coefficient, and *S* is the QS concentration required for half-maximally activate or inhibit the protein synthesis. We note that the promoter in strain Z is a hybrid one. This implies that, no matter how much of signal *s*_1_ is around, if signal *s*_2_ is sufficiently large (in particular, if *s*_2_ *≫ S*), protein synthesis in strain Z will cease. This is reflected in the model as the two regulatory Hill functions are multiply each other.

Equations (2e)-(2f) describe the synthesis rate of the quorum sensing molecules *s*_1_ and *s*_2_, respectively. The synthesis rate of each quorum sensing molecule is assumed to be proportional to the total number of synthase proteins in the corresponding strain, *i*.*e*. the synthesis rate of *s*_1_ is proportional to *N*_*x*_*x* and that of *s*_2_ is proportional to *N*_*y*_*y*, where the parameters *α*_s*i*_, *i* = 1, 2, are constants. Since in this experiment, we have not tagged the quorum sensing molecules to be degraded, we do not have a corresponding degradation term in their dynamics.

## Estimation of the Parameters of the Mathematical Model Using Experimental Data and Bayesian Inference

We used experimental data to obtain estimations of the parameters of the mathematical model. In the following, we describe what variables we measured in our experiments and how we reformulated the model to incorporate the measured variables.

### Model Reformulation to Incorporate Measured Variables

We use a plate reader that measures the total fluorescence of each strain as well as the optical density of the total cell population. We assume that the optical density of cells is proportional to the total number of cells in the well and that in our experiments we always stay in this linear range. Therefore,

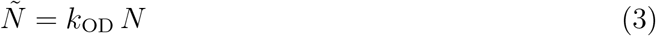

where *Ñ* is the optical density (OD), *N* is the total number of cells in the well, and *k*_OD_ is a proportionality constant.

Similarly, we assume the fluorescence signal of each strain is proportional to the total number of fluorescent proteins in that strain, *i*.*e*.

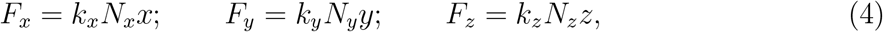

where *F*_*x*_, *F*_*y*_, and *F*_*z*_ are the fluorescence signals of strains X, Y, and Z, and *k*_*x*_, *k*_*y*_, and *k*_*z*_ are the corresponding proportionality constants.

At each experiment, we measure the time course data for *Ñ, F*_*x*_, *F*_*y*_, and *F*_*z*_. Our experiments were 8 hours (480 minutes) and we did measurements every 10 minutes.

We can rewrite Eq. (2) for the purpose of parameter inference. First, by defining the new variables 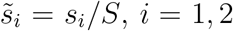, we can eliminate one parameter (*i*.*e. S*) from the model.

This is reasonable since we do not measure the concentrations of the signaling molecules *s*_1_ and *s*_2_. Second, we write the model in terms of the measured variables introduced in (3) and (4). As a result, we obtain the reformulated model

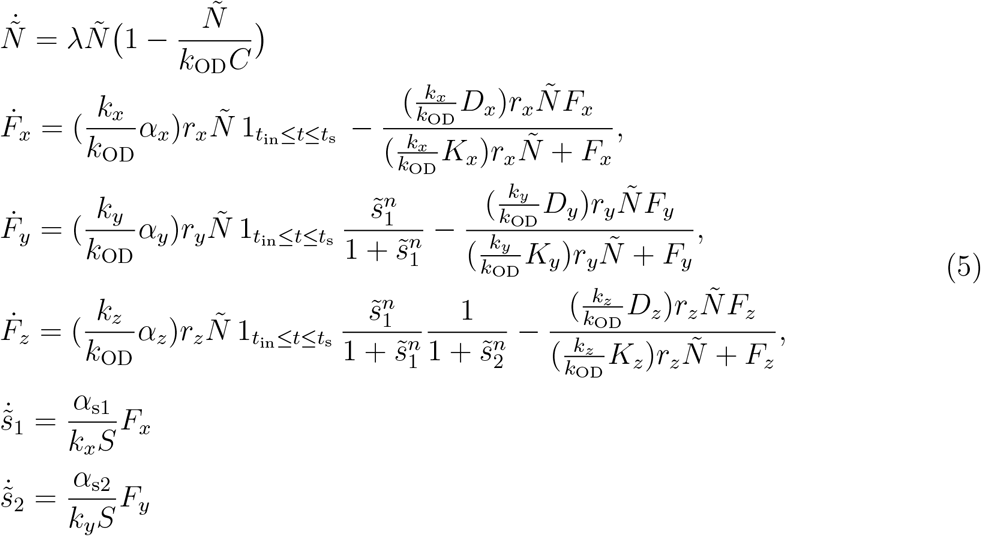

By defining the following parameters

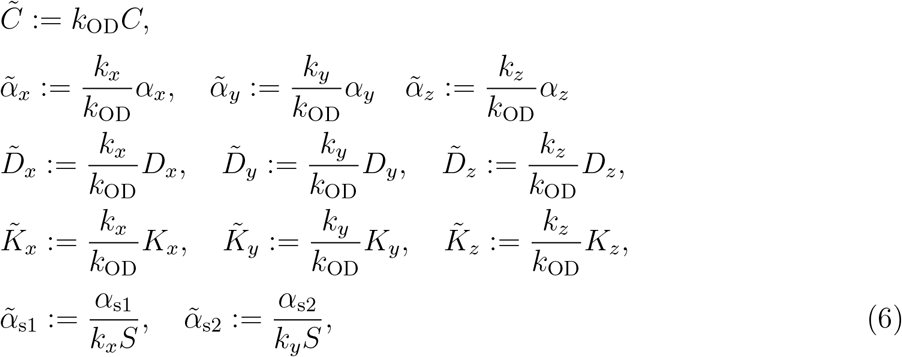

we can express model (5) more concisely as

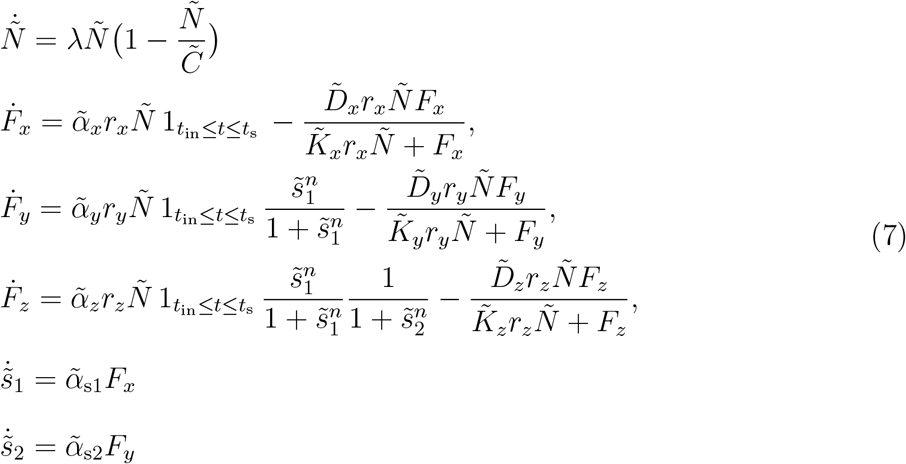

Using the Bayesian parameter inference approach outlined below, we infer the parameters in model (7). Note that most of the parameters in model (7) that are defined in (6) are not the actual physical parameters, but proportional to them. Therefore, to find the true physical values of the parameters from inference results, one needs to know the calibration constants *k*_OD_, *k*_*x*_, *k*_*y*_, and *k*_*z*_. Otherwise, we can only infer the physical parameters up to a scaling constant.

### Bayesian Parameter Inference

In Bayesian parameter inference approaches, one represents the uncertainty over the unknown parameters of the model using a prior probability density function ℙ(*θ*) where *θ* denotes a vector of unknown parameters. Then, given data from some observations or experiments, Bayes’ rule provides the posterior probability density function over the parameters, which we denote by ℙ(*θ*|data). In this setting, ℙ(*θ*) is called the prior probability distribution as it represents our prior knowledge about parameters. The Bayes’ rule states that

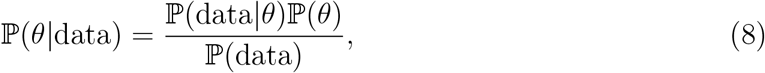

where ℙ(data|*θ*) is called the likelihood function and ℙ(data) is the integration of the likelihood over the space of parameters. Given the data, ℙ(data) becomes a constant and therefore, the posterior distribution is proportional to the product of the likelihood and prior distribution.

The goal is to find the posterior distribution ℙ(*θ*|data), however, in general this distribution is difficult to obtain directly. Therefore, it is often necessary to approximate the posterior distribution using a number of samples from the right hand side of (8). From these samples, we can then obtain different parameter distributions of interest, in particular, marginal distributions or pairwise distributions. One may then estimate parameters using the maximum (MAP estimate) or mean of these approximate posterior distributions.

Markov Chain Monte Carlo (MCMC) techniques are often used to obtain samples from the posterior distribution. The sequence of sampled parameters form a Markov chain and it can be shown that the limiting probability distribution of this Markov chain is the posterior distribution of interest. Below we describe the details of our Bayesian method.

### Data

For the data we use 3 experiments conducted at the target fractions (*r*_*x*_ = 0.4667, *r*_*y*_ = 0.2), (*r*_*x*_ = 0.2, *r*_*y*_ = 0.4667) and (*r*_*x*_ = 0.2, *r*_*y*_ = 0.2). Each experiment contains the timecourse data of the variables *Ñ, F*_*x*_, *F*_*y*_, and *F*_*z*_, that are respectively, the OD and the fluorescence signals of strains X, Y, and Z.

We also tested using only the data for one experiment, in particular, the experiment conducted at (*r*_*x*_ = 0.2, *r*_*y*_ = 0.2). In this case, the prediction capability of the model slightly deteriorated, though still can be satisfactory (see Appendix Fig. S9).

### Model and Parameters

We used model (7) which has a total of 14 unknown parameters. Since our circuit is an induced system, the initial values of all fluorescence signals and the signalling molecules were assumed to be zero. The initial OD was consistent in our experiments and based on the data we set it to *Ñ*_0_ = 0.008577. We performed a preliminary sensitivity analysis of the model, and fit the model to test data (separate from the data we used for inference). This analysis showed that some model parameters are either difficult to identify jointly, or the model is insensitive to their actual values. For these parameters we used values that were broadly consistent with values found in our preliminary fits, and are consistent with values found in the literature, when available. These parameters were the Michaelis-Menten constants and the Hill coefficient of the regulatory terms. Therefore, based on our preliminary investigations, we set the values of these parameters to: 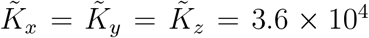 and *n* = 2. Moreover, the induction time was set at *t*_in_ = 120 min in all experiments. We approximated the start of the stationary phase of the cell growth was set to *t*_s_ = 270 min (which is approximated from the data). The set of 10 remaining unknown parameters are therefore

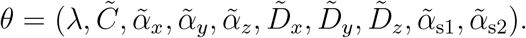

### Error Model and Likelihood Function

We assumed that there were measurement errors for OD and the fluorescence signals, and that these measurement errors were indpendent and followed a normal distribution with mean zero. We found the standard deviations of the OD and the fluorescence measurement errors using data from separate experiments: 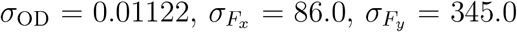, and 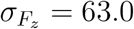.

The likelihood function, given the assumptions above, is

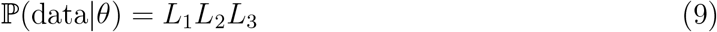

where *L*_1_, *L*_2_, and *L*_3_ refer to the likelihood terms corresponding to the three experiments we used for the inference. For each *e, e* = 1, 2, 3, we have

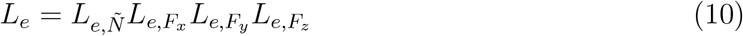

where

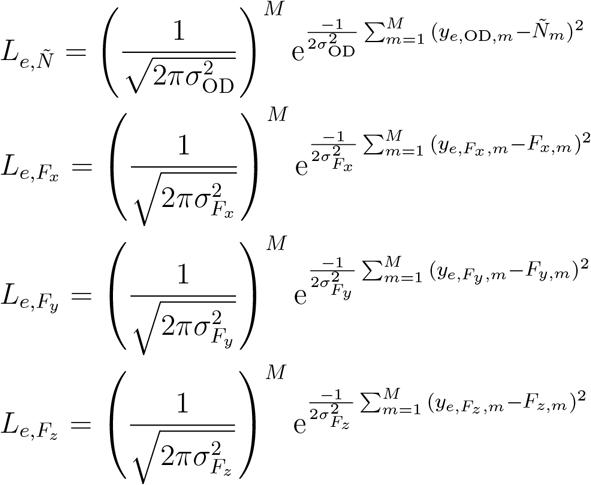

where *m, m* = 1, …, *M* is an index for the data time points (*M* = 49 in our data), 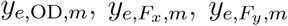, and 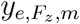 are the data associated with experiment *e* and for the OD and the fluorescence signals of strains X, Y, and Z, respectively, and *Ñ*_*m*_, *F*_*x,m*_, *F*_*y,m*_, and *F*_*z,m*_ are the model values generated from a simulation of model (7) for the given parameters *θ*. Since the model values depend on the parameters *θ*, the likelihood also depends on *θ*.

### Priors

We chose some of our experiments for preliminary investigations in order to find appropriate, narrower priors for the parameters. At first for each parameter, we chose a uniform prior distribution over a relatively big interval and fit the model to data from these preliminary experiments. Then, used the resulting posterior distribution to narrow the priors in the actual fits.

Furthermore, the initial investigations showed that the parameters have different ranges and scales. In general, the MCMC algorithms work more efficiently if the parameters’ orders of magnitude are similar. Hence, we rescaled some of the parameters as shown below

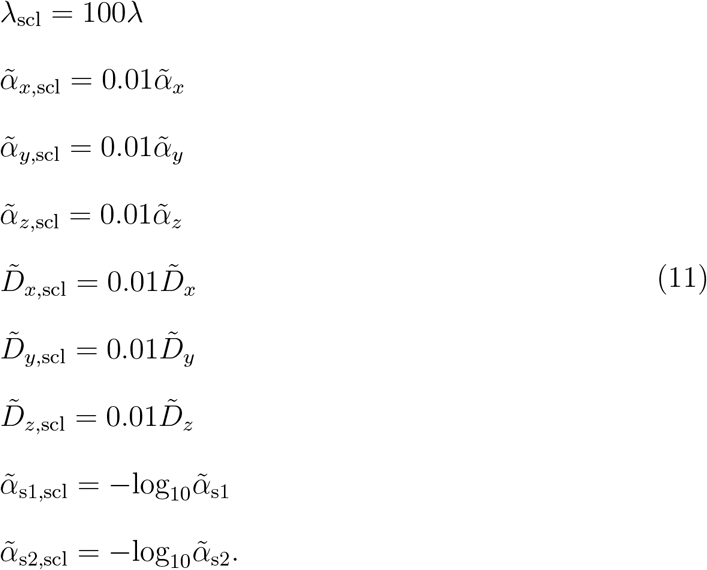

After the rescaling, we chose the following priors for the parameters

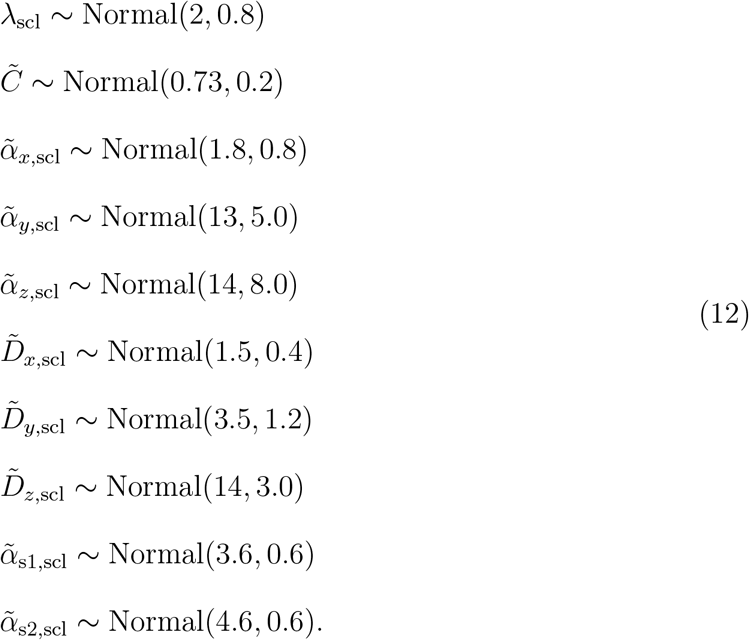

where Normal(*a, b*) denotes the normal probability distribution with mean *a* and standard deviation *b*.

### Hamiltonian Markov Chain Monte Carlo

Once one has the data, model, unknown parameters, error model, and priors, one can perform MCMC techniques to sample the posterior distribution. We used the software package Stan (https://mc-stan.org/) that implements the state-of-the-art Hamiltonian Monte Carlo method. Details about our implementation of this method can be found at the Github page which contains our code and data.

https://github.com/biodesign-lab/Predictable_tuning_2020

### Results of Model Fitting to Data

After we performed sampling, we obtained samples from the 10-dimensional posterior distribution. One way to visualize the posterior distribution is to look at the marginal distribution of each parameter as well as the pairwise distributions of every pair of parameters (Appendix Fig. S11). The panels on the diagonal of Appendix Fig. S11 show the marginal distribution, while the other panels show the pairwise distributions. The parameters for which the posterior distribution is maximal (MAP estimates) are given below. We used these parameters for model prediction. Appendix Fig. S10 shows the fit of the model (using the MAP estimate shown in Table 1) to the experimental data.

**Table 1:** The MAP estimate and the credible intervals for the parameters

| Scaled parameters | %95 Credible interval |  | Parameters of model (7) | %95 Credible interval |
| --- | --- | --- | --- | --- |
| $\lambda_{\text{scl,MAP}} = 2.05$ | $2.04 \leq \lambda_{\text{scl}} \leq 2.06$ | | $\lambda_{\text{MAP}} = 0.0205$ | $0.0204 \leq \lambda \leq 0.0206$ |
| $\tilde{C}_{\text{MAP}} = 0.738$ | $0.734 \leq \tilde{C} \leq 0.741$ | | $\tilde{C}_{\text{MAP}} = 0.738$ | $0.734 \leq \tilde{C} \leq 0.741$ |
| $\tilde{\alpha}_{x,\text{scl,MAP}} = 1.85$ | $1.81 \leq \tilde{\alpha}_{x,\text{scl}} \leq 1.94$ | | $\tilde{\alpha}_{x,\text{MAP}} = 185$ | $181 \leq \tilde{\alpha}_x \leq 194$ |
| $\tilde{\alpha}_{y,\text{scl,MAP}} = 13.27$ | $13.05 \leq \tilde{\alpha}_{y,\text{scl}} \leq 13.53$ | | $\tilde{\alpha}_{y,\text{MAP}} = 1327$ | $1305 \leq \tilde{\alpha}_y \leq 1353$ |
| $\tilde{\alpha}_{z,\text{scl,MAP}} = 14.05$ | $13.68 \leq \tilde{\alpha}_{z,\text{scl}} \leq 14.65$ | | $\tilde{\alpha}_{z,\text{MAP}} = 1405$ | $1368 \leq \tilde{\alpha}_z \leq 1465$ |
| $\tilde{D}_{x,\text{scl,MAP}} = 1.52$ | $1.44 \leq \tilde{D}_{x,\text{scl}} \leq 1.68$ | Scaling back<br>⇒ | $\tilde{D}_{x,\text{MAP}} = 152$ | $144 \leq \tilde{D}_x \leq 168$ |
| $\tilde{D}_{y,\text{scl,MAP}} = 3.26$ | $3.15 \leq \tilde{D}_{y,\text{scl}} \leq 3.34$ | | $\tilde{D}_{y,\text{MAP}} = 326$ | $315 \leq \tilde{D}_y \leq 334$ |
| $\tilde{D}_{z,\text{scl,MAP}} = 13.94$ | $13.26 \leq \tilde{D}_{z,\text{scl}} \leq 14.45$ | | $\tilde{D}_{z,\text{MAP}} = 1394$ | $1326 \leq \tilde{D}_z \leq 1445$ |
| $\tilde{\alpha}_{s1,\text{scl,MAP}} = 3.593$ | $3.577 \leq \tilde{\alpha}_{s1,\text{scl}} \leq 3.628$ | | $\tilde{\alpha}_{s1,\text{MAP}} = 2.55 \times 10^{-4}$ | $2.36 \times 10^{-4} \leq \tilde{\alpha}_{s1} \leq 2.65 \times 10^{-4}$ |
| $\tilde{\alpha}_{s2,\text{scl,MAP}} = 4.642$ | $4.597 \leq \tilde{\alpha}_{s2,\text{scl}} \leq 4.668$ | | $\tilde{\alpha}_{s2,\text{MAP}} = 2.28 \times 10^{-5}$ | $2.15 \times 10^{-5} \leq \tilde{\alpha}_{s2} \leq 2.53 \times 10^{-5}$ |

Code and data for our Bayesian inference analysis and results, as well as code for generating all figures can be found in our Github repository.

https://github.com/biodesign-lab/Predictable_tuning_2020

## Author Contributions

D.M.Z., M.S., W.O., K.J., and M.R.B. designed the study; D.M.Z. and S.M. performed the experiments; R.N.A. and A.J.H. designed and fabricated the microfluidic device; M.S., W.O., and K.J. designed and implemented the computational model; D.M.Z., M.S., W.O., K.J., and M.R.B. analyzed the results; Every author wrote the paper.

## Notes

The authors declare that they have no conflict of interest.

## Acknowledgement

This work was funded by the National Institutes of Health and the National Sciences Foundation through the joint NSF-National Institutes of General Medical Sciences Mathematical Biology Program grant nos. DMS-1662290 (M.R.B.) and DMS-1662305 (K.J.); NSF grants MCB-1936770 (K.J.), MCB-1936774 (M.R.B.), and DMS-1816315 (W.O.); NSF Graduate Research Fellowship Program grants GRFP-1450681 (D.M.Z.) and GRFP-1842494 (R.N.A.); the National Institutes of Health grant no. R01GM117138 (M.R.B., K.J., W.O.); the Robert A. Welch Foundation grant no. C-1729 (M.R.B.); and the Hamill Foundation (M.R.B.). The authors thank Andrea Padrón for graphics of lab equipment used in the figures. The authors acknowledge the use of resources of the Shared Equipment Authority at Rice University for this work.

## Supporting Information Available

The following files are available free of charge.

- Plasmids used in this study (Table S1)
- Schematic of the microfluidic device (Figure S1)
- Fluorescent microscopy experiments of the consortial feedforward loop (Figure S2)
- Control circuits known not to pulse (Figure S3)
- Additional replicates (Figure S4)
- Barcoding and NGS strategy (Figure S5)
- Target vs Measured strain fractions (Figure S6)
- Additional replicates (Figure S7)
- Strain fractions at the start and end of experiment (Figure S8)
- Model predictions based on 1 vs 3 experiments (Figure S9)
- Simulations of the model against experimental data (Figure S10)
- Posterior Distributions (Figure S11)
- Target and measured fractions for same amplitude experiments (Figure S12)
- Plating assay (Figure S13)

## TOC Graphic

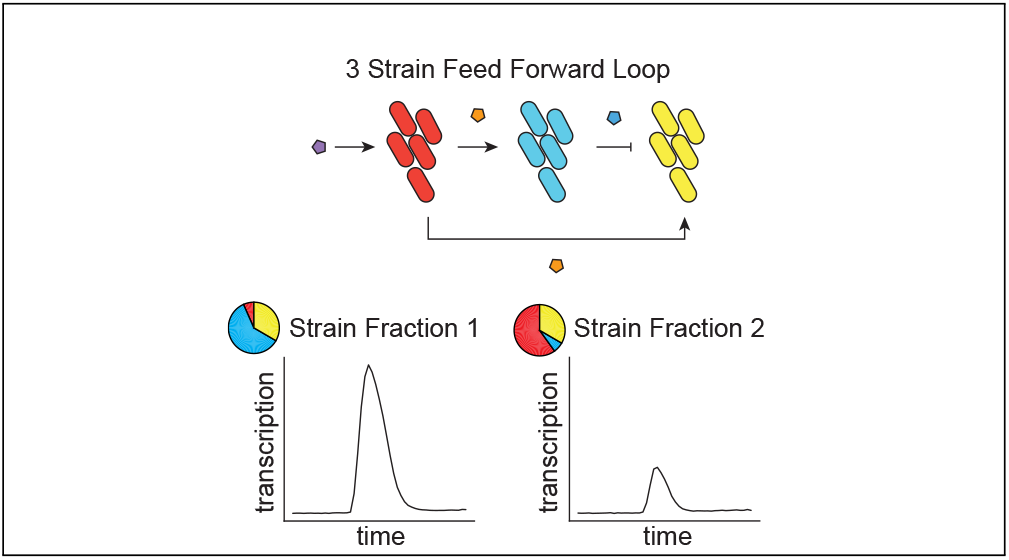

