## Supplementary file for "Tunable dynamics in a multi-strain transcriptional pulse generator"

### Supplementary Information - Tunable dynamics in a multi-strain transcriptional pulse generator

Table S1: Plasmids used in this study.

| Plasmid | Cassettes | Origin | Resistance | Used in Strain |
|---|---|---|---|---|
| pDZ033 | $P_{Cin}$ *rhlI-ssrA*, $P_{Cin}$ *sfCFP-ssrA* | ColE1 | $Kan^R$ | Y |
| pDZ047 | Blank Plasmid | p15A | $Amp^R$ | X, Y, Z Cascade, Z Fanout |
| pDZ113 | $P_{Rhl}$ *sfYFP-ssrA* | ColE1 | $Kan^R$ | Z Cascade |
| pDZ114 | $P_{CinLac}$ *sfYFP-ssrA* | ColE1 | $Kan^R$ | Z, Z Fanout |
| pDZ115 | $P_{LLacO1}$ *sfYFP-ssrA* | ColE1 | $Kan^R$ | Z Inverter |
| pDZ134 | $P_{Rhl}$ *rbsR-L-ssrA* | p15A | $Amp^R$ | Z, Z Inverter |
| pDZ149 | $P_{LLacO1}$ *cinI-ssrA*, $P_{LLacO1}$ *mCherry2-ssrA* | ColE1 | $Kan^R$ | X |

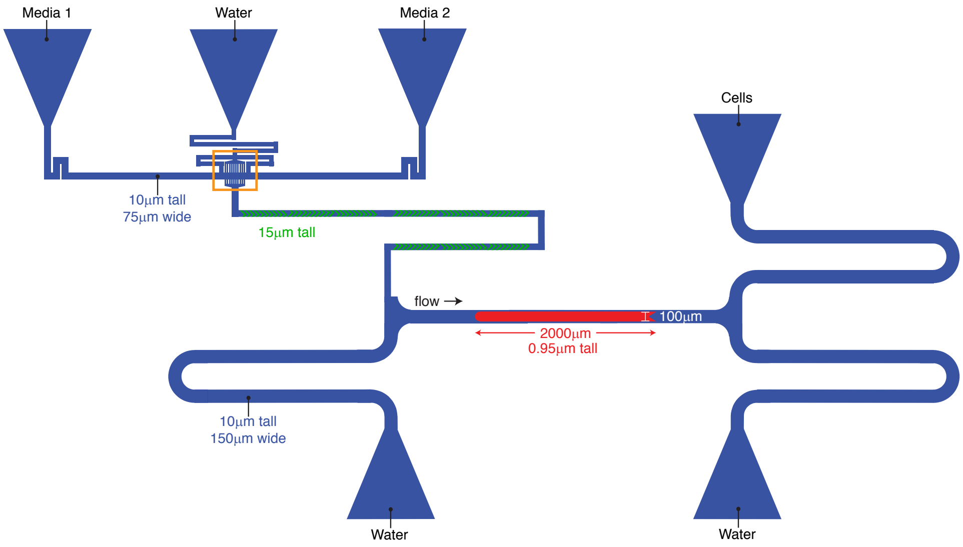

Figure S1: Schematic of the microfluidic device. Blue regions are the flow channels and the red region is the cell-trapping region. Blue triangles are ports to which media, cells, and water waste reservoirs are connected. The 'dial-a-wave' junction boxed in orange between the two media input ports allows for mixing different ratios of the two media types. Green mixers following the 'dial-a-wave' junction are raised regions of the flow channel to allow for proper mixing of the two media types.

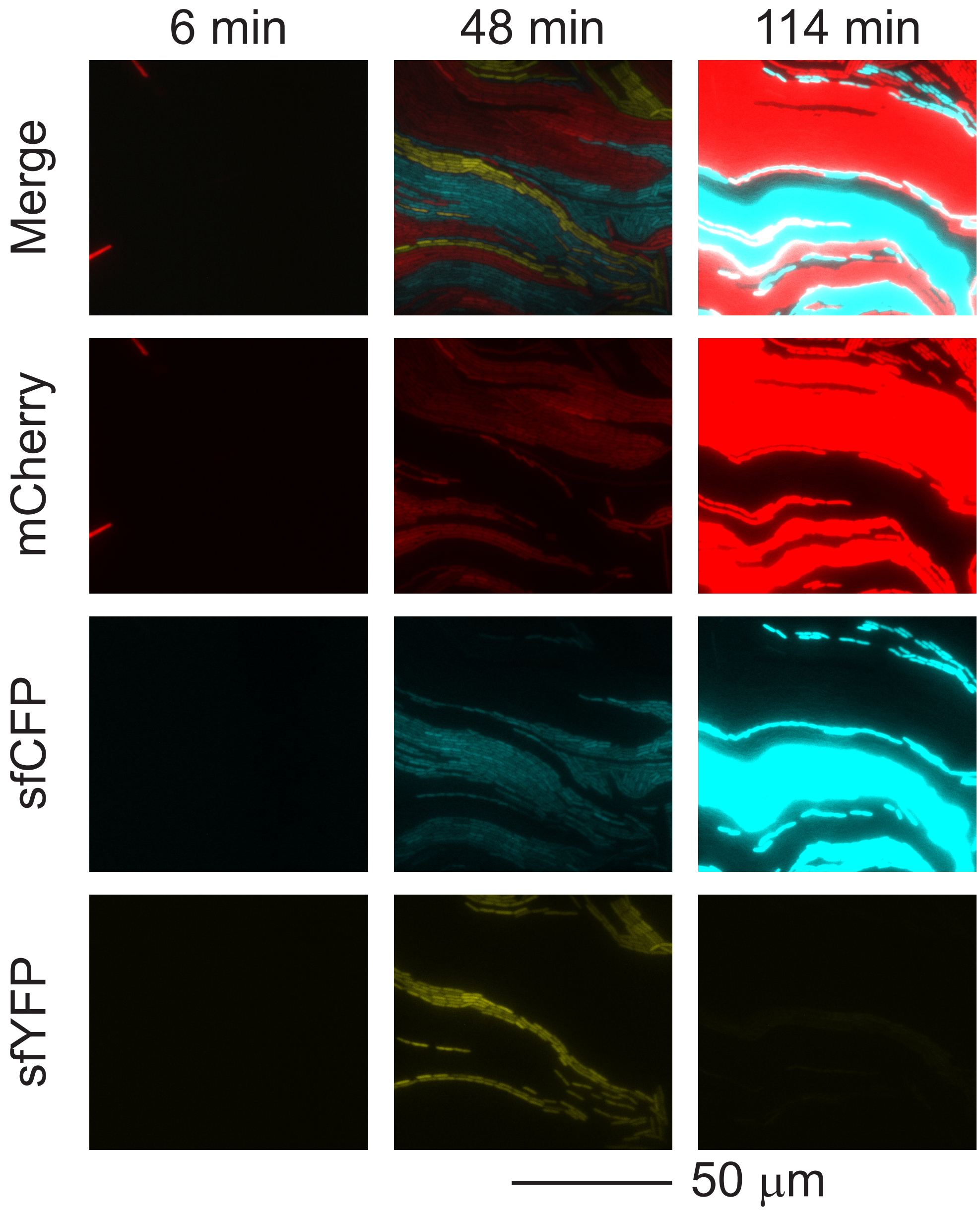

Figure S2: Fluorescence microscopy experiments of the consortial feedforward loop. 2 mM IPTG was introduced to the cell trapping region at time=0 minutes. Images show the same portion of the microfluidic device at 6, 48, and 114 minutes. Note that the sfYFP pulses in time.

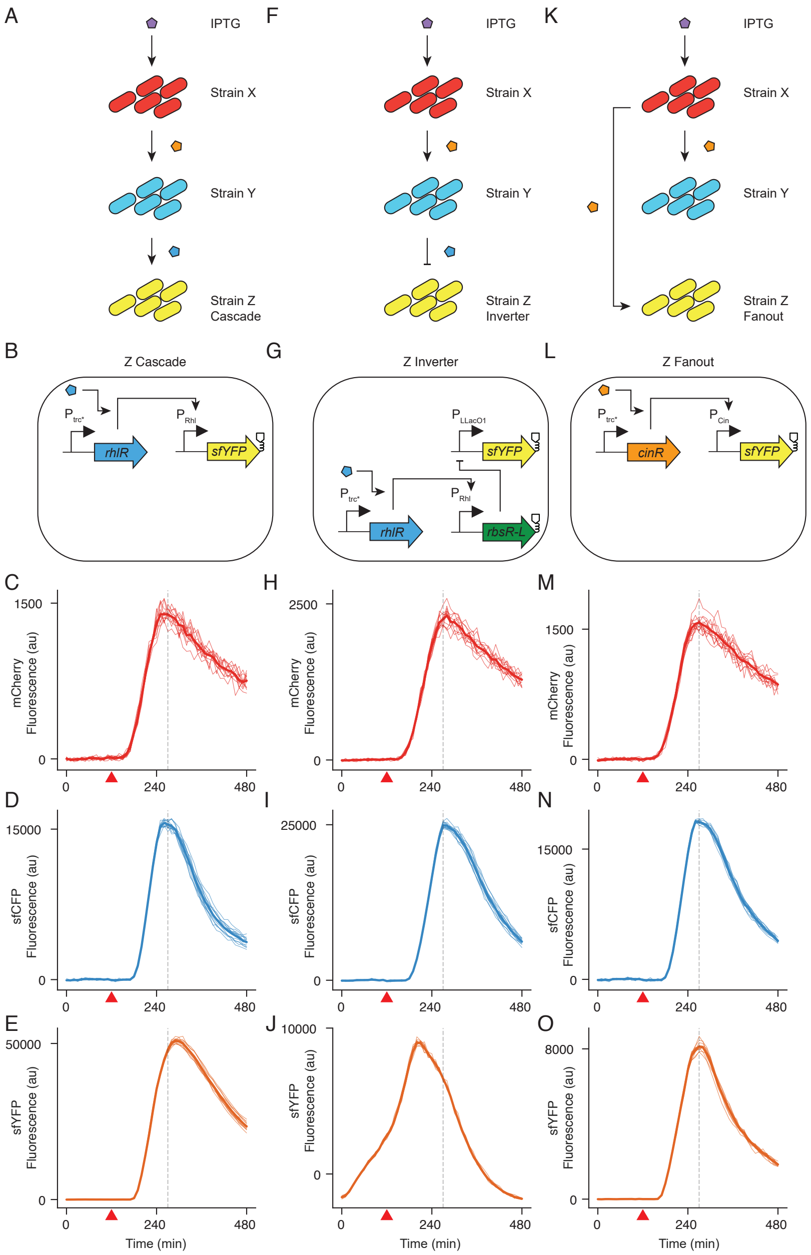

Figure S3: Control circuits known not to pulse. In all three control circuit experiments, strain Z was replaced with a version of strain Z that changes the circuit topology. (*A*) Circuit topology of the cascade circuit. (*B*) Gene circuit for the strain Z cascade. (*C,D,E*) Timeseries result for the cascade circuit. Note that in (*E*), sfYFP does not pulse before the onset of stationary phase. (*F*) Circuit topology of the inverter circuit. (*G*) Gene circuit for the strain Z inverter. (*H,I,J*) Timeseries result for the inverter circuit. Note that sfYFP signal begins to decline before the onset of stationary phase. (*K*) Circuit topology of the fanout circuit. (*L*) Gene circuit for the strain Z fanout. (*M,N,O*) Timeseries result for the fanout circuit. As with the cascade circuit, sfYFP does not pulse before the onset of stationary phase. Red triangle indicates the time at which IPTG was added (120 minutes).

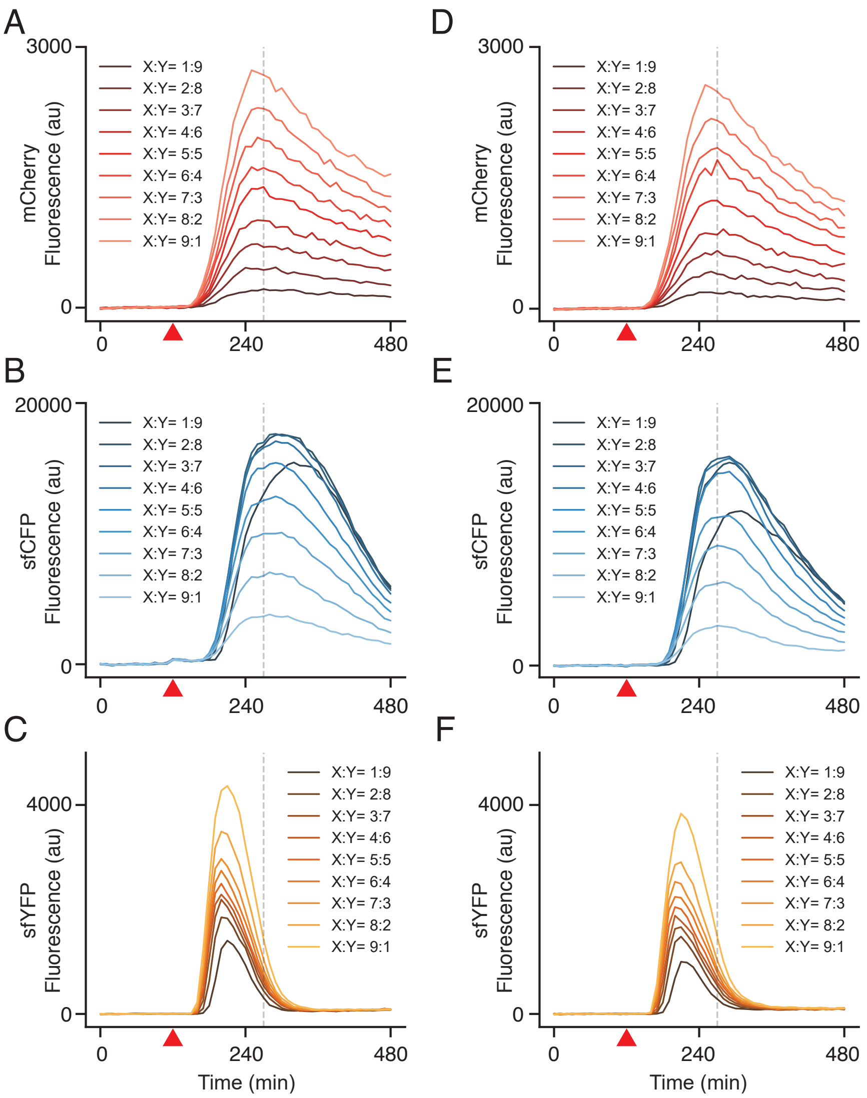

Figure S4: Additional replicates of the experiment where Z was fixed to a population fraction of 33%. ($A$,$B$,$C$) Fluorescence of strains X, Y, and Z, respectively, for one day of experiments. Each curve represents the mean of 6 technical replicates. ($D$,$E$,$F$) Another replicate of the same experiment, performed on a different day. Red triangle indicates the time at which IPTG was added (120 minutes).

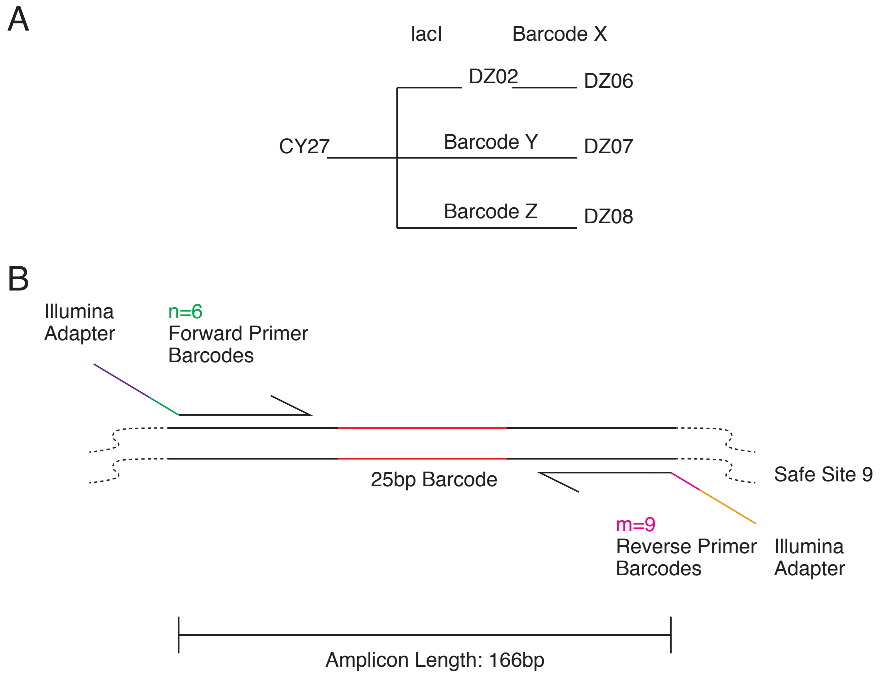

Figure S5: Barcoding and next-gen sequencing strategy. (*A*) base strain CY27 was modified to have constitutively expressed lacI (DZ02) and further modified with a unique 25bp barcode (DZ06). CY27 was modified with unique barcodes to represent strain Y and Z (DZ07 and DZ08 respectively). (*B*) PCR amplification of barcode from DZ06,07,08 genome. Unique barcode was captured in amplicons using PCR primers that bound to all three strains. Primers contained Illumina adapters on the 5' end, followed by a 5 bp index. A different forward primer reverse primer pair was used for each PCR reaction then resulting reactions were pooled together and sent for sequencing. Results were demultiplexed using the index sequences and strain fraction was determined by counting occurrences of X, Y, and Z barcode sequences.

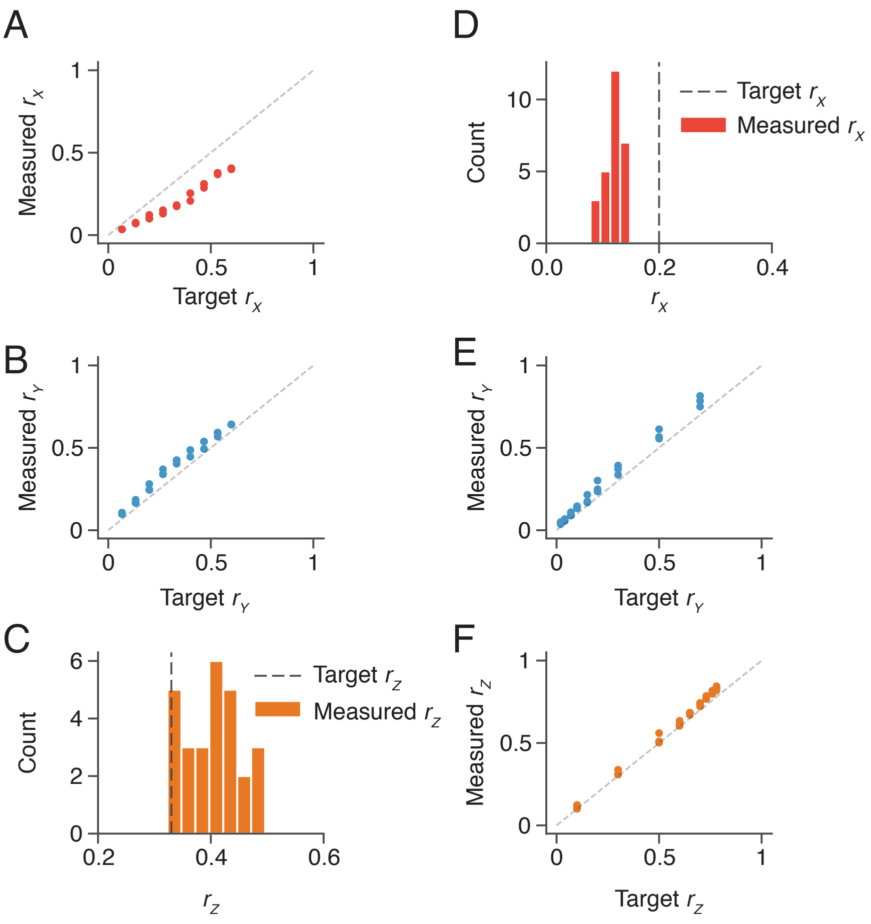

Figure S6: Target strain fractions versus measured strain fractions at the onset of the experiment. ($A$,$B$,$C$) Target strain fractions versus measured strain fractions from the experiment where strain Z was set to 33% and strains X and Y were varied. ($D$,$E$,$F$) Target strain fractions versus measured strain fractions from the experiment where strain X was set to 20% and strains Y and Z were varied.

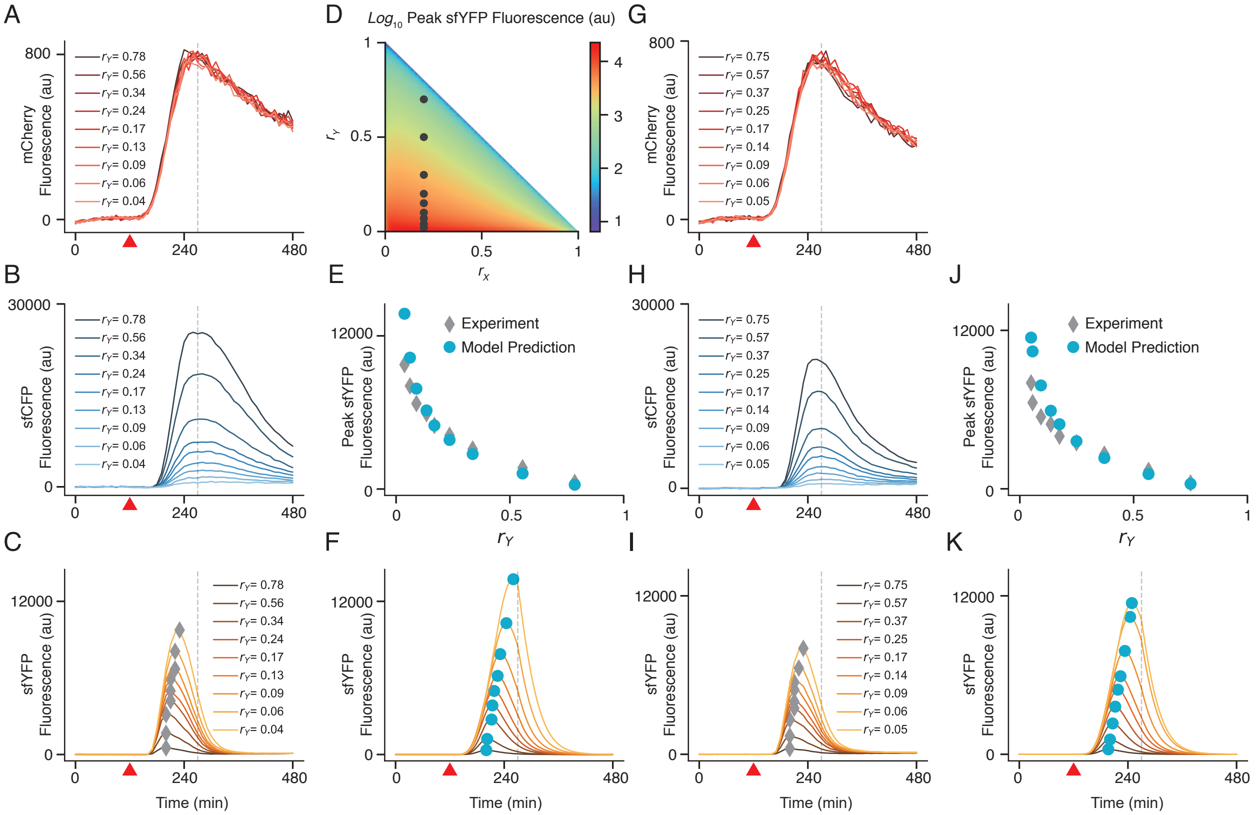

Figure S7: Additional replicates of the experiment where X was fixed to a population fraction of 20%. (*A,B,C*) Experimental results showing signal of strains X, Y, and Z, respectively, with strain fractions for strain Y as indicated. Gray diamonds indicate maximal fluorescence. (*D*) Target fractions shown as black circles in fraction space. (*E*) Model prediction for peak sfYFP height (blue circles), compared with experimental data (gray diamonds). (*F*) Timeseries from simulation of strain Z activity. Blue circles indicate peak fluorescence. (*G,H,I*) Repeat of the same experiment performed on a different day. (*J*) Comparison of strain Z activity to model prediction. (*K*) Simulation of strain Z activity. Red triangle indicates the time at which IPTG was added (120 minutes).

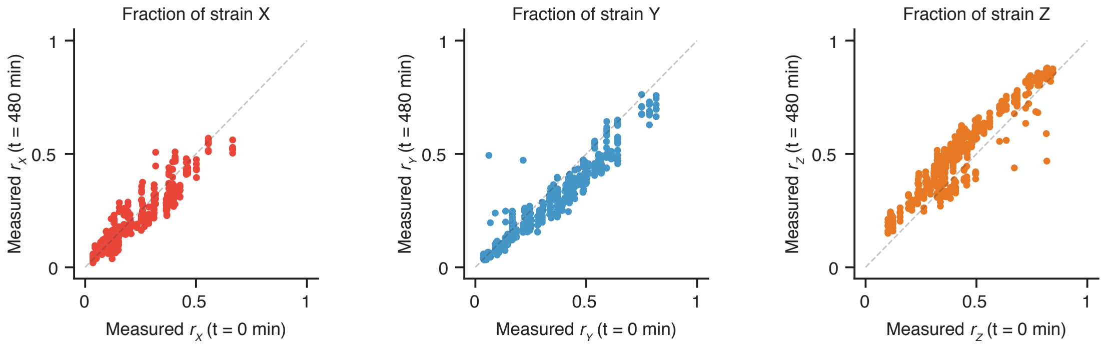

Figure S8: Strain fractions measured using next-gen sequencing before the start of and after the end of each experiment performed. Plots show measured strain fractions across all experiments performed.

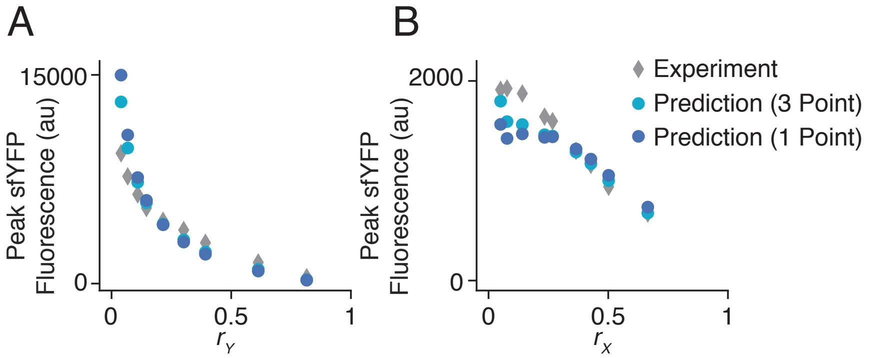

Figure S9: Comparison of model predictions based on 1 experiment (1 strain fraction point) to those based on 3 experiments (3 strain fraction points). (*A*) Predictions for experiment where strain X is fixed to 20%. (*B*) Predictions for experiment where peak heights were meant to be the same .

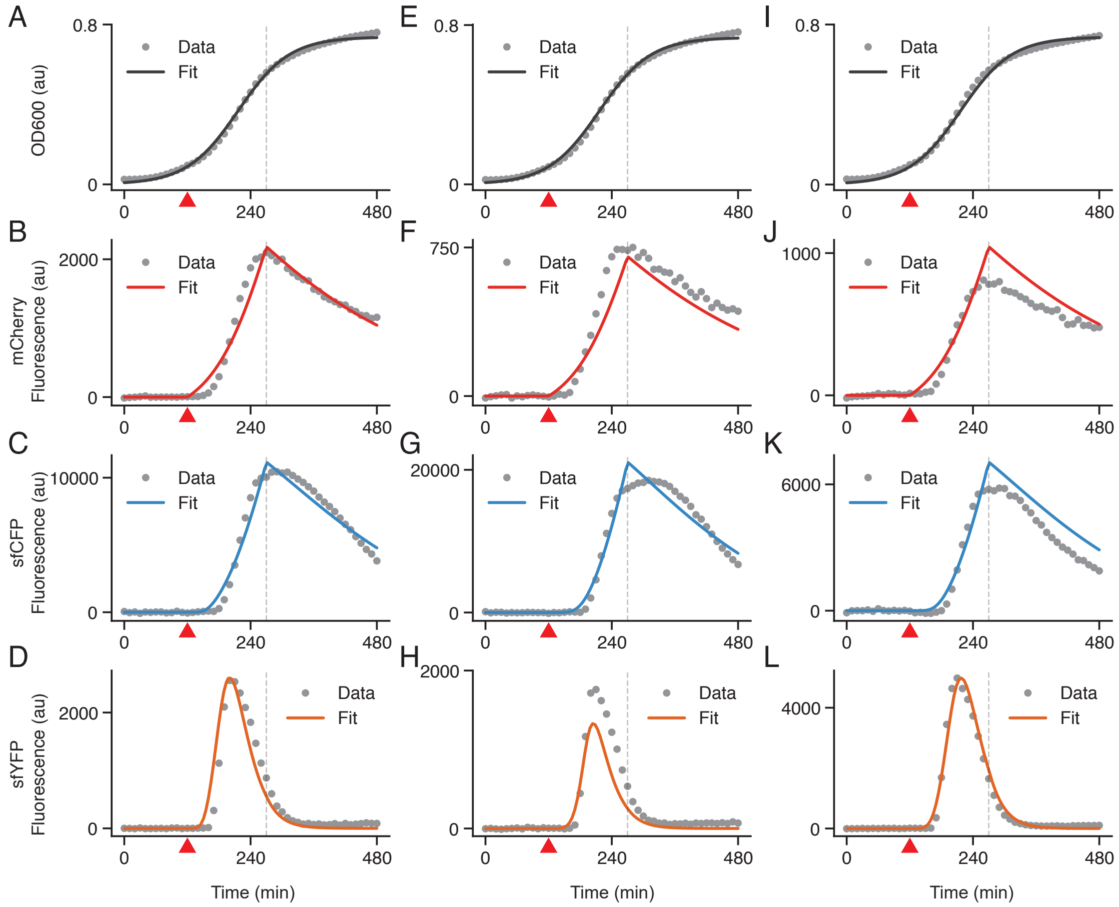

Figure S10: Simulations of the mathematical model (solid curves) against experimental data (gray dots). For model simulation, the fitted parameters (MAP estimate) are used. The first column (*A-D*) compares the simulation of the mathematical model to the data from the first experiment conducted at $(r_x = 0.47, r_y = 0.2)$. The second column (*E-H*) and the third column (*I-L*) show the same for the experiments conducted at $(r_x = 0.2, r_y = 0.47)$ and $(r_x = 0.2, r_y = 0.2)$, respectively. Red triangle indicates the time at which IPTG was added (120 minutes).

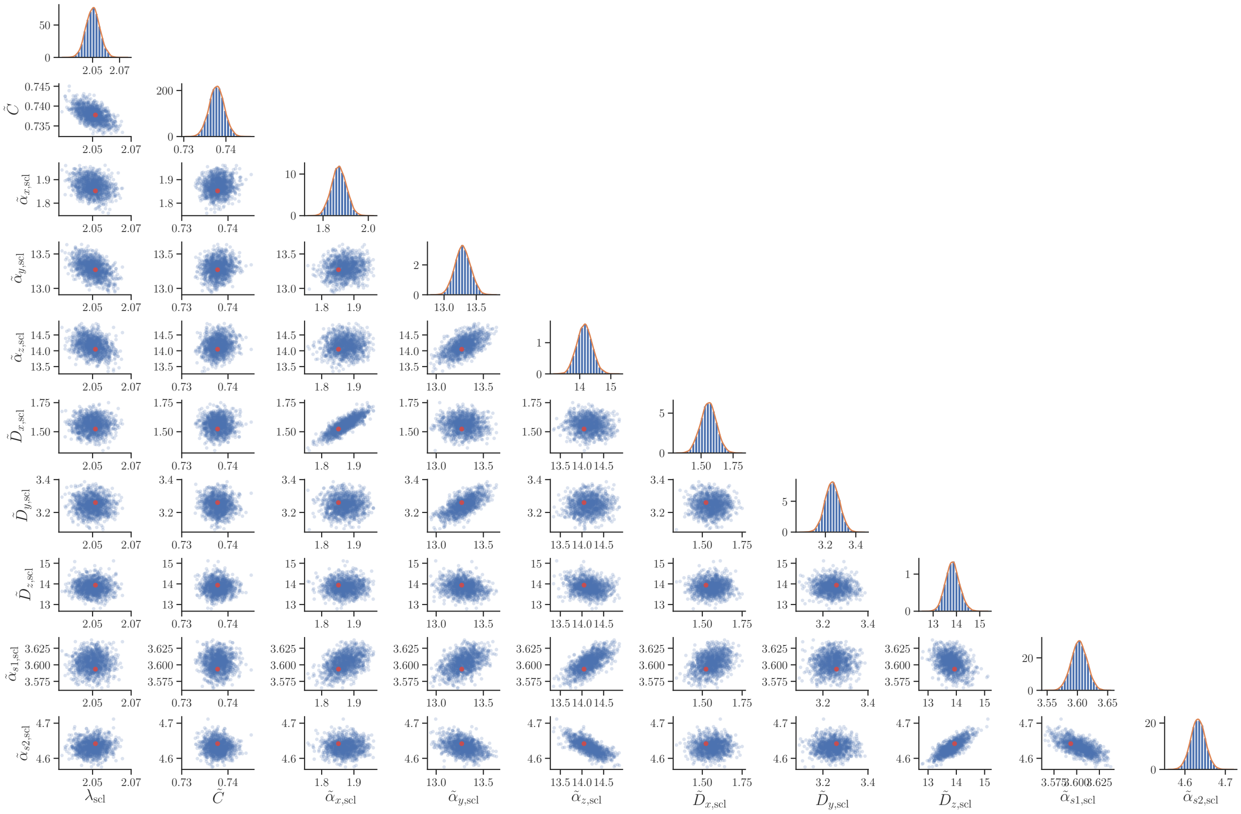

Figure S11: Posterior distributions. The panels on the diagonal show the marginal posterior distribution for each parameter. The panels below diagonal show the pairwise posterior distributions for each pair of parameters. The red dot indicates the MAP estimate.

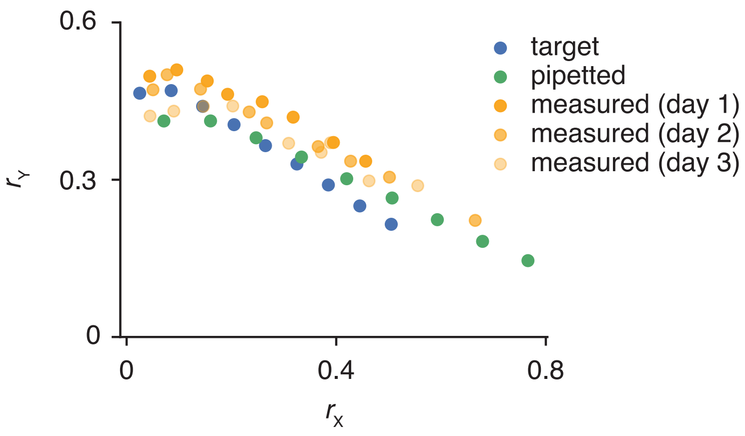

Figure S12: Target and actual fractions for same peak height experiment. Using data from previous experiments, we calculated a linear transformation of the target $r_x, r_y$ to pipette. Resulting measured fractions appeared to be closer to the target fractions than the pipetted fractions.

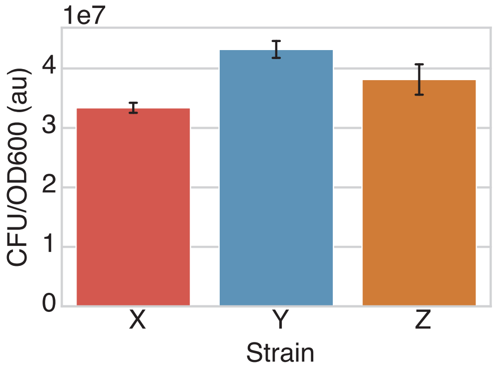

Figure S13: Colony forming units (CFU) per unit OD600 for each strain as measured using a plating assay. Bars are the mean of 5 replicates performed on different days. Error bars represent the standard deviation. CFU/OD600 results are consistent with next-gen sequencing data because strain X is consistently lower than strains Y and Z.
